# Non-senescent species are not immortal: stress and decline in two planaria species

**DOI:** 10.1101/2023.01.12.523873

**Authors:** Jacques A. Deere, Penelope Holland, Aziz Aboobaker, Roberto Salguero-Gomez

**Author notes:** Corresponding authors: Institute for Biodiversity and Ecosystem Dynamics (IBED), University of Amsterdam, P.O. Box 94249, 1090 GE Amsterdam, The Netherlands Department of Biology, University of Oxford, South Parks Road, OX1 3RB, Oxford, UK.

## Abstract

1. Potential immortality is observed in several species (*e.g.*, prickly pear cactus, hydra, flatworms) and is indicative of their negligible or even negative senescence rates. Unlike in senescent species, which experience reduced individual performance with age due to physiological degradation, species with negligible or negative senescence display mortality rates that remain constant or decline with age, respectively. These rates vary across taxa and are correlated with life history traits. Yet, the extent to which variable resource availability, a key driver of variation in life history traits, impacts species that show negligible or negative senescence is currently unknown.

2. Here, we examine whether and how variation in the quantity, quality, and feeding interval of resources impact population structure, population performance, and life history trait trade-offs in two long-lived planaria that do not senesce: *Schmidtea mediterranea* and *Dugesia tahitiensis*. In a full factorial design, different combinations of resource quantity (reduced intake, standard intake and high intake) and quality (high and low quality) were provided in two different feeding intervals (7-day and 14-day intervals) for 19 weeks.

3. We show that variability in resource availability, via decreases in quantity, quality, and frequency of resources, does not diminish population viability in either species but does result in suboptimal conditions of stress in *S. mediterranea*.

4. The high population viability we report can be attributed to two different mechanisms: increased reproduction or increased investment into maintenance at the expense of reproduction. Moreover, which mechanism was responsible for said high population viability was context-dependent and modulated by the specific life history strategy of the two planaria species.

5. We show that suboptimal conditions can cause stress responses that have significant impacts on non-senescent species. The context-dependent response we observe suggests that species that do not senesce but are subject to suboptimal conditions of stress may ultimately exhibit declines in performance and ultimately die. A clearer understanding of the impact of suboptimal conditions of resource availability on non-senescent species is needed to determine the extent of stress experienced and ultimately whether a species can truly be immortal.

## INTRODUCTION

A hydra (a potentially immortal metazoan (Martínez, 1998)) walks out of a bar… and is run over by a bus. Nothing lasts forever, including potential immortality. How age impacts individual performance and fitness has received much attention (Baudisch, 2011; Baudisch & Stott, 2019; Caswell & Salguero-Gómez, 2013; Kirkwood, 1977; Medvedev, 1990; J. W. Vaupel, 2010). The reduction of individual performance with age due to physiological degradation, senescence (Finch, 1994; Jones et al., 2008), was initially thought to be universal, although recent evidence shows that not all species reduce individual performance with age (review by Flatt & Partridge, 2018; (Caswell & Salguero-Gómez, 2013; Jones et al., 2008, 2014; Roper et al., 2021; but see Nelson & Masel, 2017). Indeed, some species like the ocean quahog (*Arctica islandica*) (Abele et al., 2009), sugar cone pine (*Pinus lamertiana*) or guppy (*Poecilia reticulata*) (Roper et al., 2021) have mortality rates that remain constant with age, while other species display mortality rates that decline with age (Roper et al., 2021; Vaupel et al., 2004). The former phenomenon is known as negligible senescence (Finch, 1994, 2009; Vaupel et al., 2004) and the latter as negative senescence (Jones et al., 2014; Vaupel et al., 2004). Negative senescence has been reported in hydra (Martínez, 1998), prickly pear cactus (*Opuntia rastrera*) (Roper et al., 2021), or flatworms (planaria, Phylum Platyhelminthes) (Sahu et al., 2017). Crucially, regardless whether a species is senescent, negligibly senescent, or negatively senescent, its rates of senescence typically correlate with other life history traits (*e.g.*, generation time; Healy et al., 2019; Salguero-Gómez et al., 2016), with the magnitude of actuarial (= survival) senescence largely coupled by the speed of the life history (Jones 2008).

The underlying correlation between life history traits and rates of senescence then raises the question of how drivers of variation in life history traits ultimately impact senescence. Resource availability is a key driver of individual development and population performance (Ozgul et al., 2009; Smallegange, 2011; Stearns, 1992). Yet, studies on how resource availability impacts the life cycle of an organism have largely focused on specific life transitions (*e.g.*, age at maturity (Auer, 2010; Smallegange, 2011; Stearns, 1992) or features of the life cycle (*e.g.*, reproduction (Rutstein et al., 2005; Williams, 1996); survival (Ozgul et al., 2009)). Survival, development, and reproduction are key demographic processes in the life cycle of any species. Indeed, these demographic processes define life history traits (*e.g.*, age at maturity, longevity, etc.) that, in turn, produce life history strategies that shape individual fitness and dynamics of a population (Roff 1992; Stearns 1992; Capdevila & Salguero-Gómez 2019). Life history theory predicts a trade-off in resource allocation between maintenance (survival and development) and reproduction of individuals (Roff, 1992; Stearns, 1992). However, the impact of resource availability on these demographic processes, and therefore on life history traits, is not straightforward. The impacts of variability in resources on demographic process can be non-linear (*e.g.*, indirect winter climate on summer foraging conditions; Mysterud et al., 2001) but can also interact with other factors (*e.g.*, anthropogenic land use change) resulting in demographic impacts being exacerbated or buffered (Maron et al., 2015).

Given the nuanced impact that resources have on life history traits, a key question in ecology and evolution is to what extent negligibly and negatively senescent species are impacted by variable resource availability. Here, we contribute to filling this gap of knowledge by examining the impact of variability of resources in two long-lived invertebrate species, the planarians *Schmidtea mediterranea* (Benazzi, Baguna, Ballister, Puccinelli & Del Papa, 1975) and *Dugesia tahitiensis* (Gourbault, 1977). Obligate asexual planaria do not show physiological senescence (Sahu et al., 2017). In these animals, stem cells also function as germ line, and so many of the well-known somatic mechanisms of senescence (Finkel & Holbrook, 2000; Lemaître et al., 2024; Payne & Chinnery, 2015) do not apply to obligate asexual planarian stem cells (Sahu et al., 2017). However, individuals of planarians can still suffer suboptimal conditions of stress, shrink, and even die (*e.g.*, Ofoegbu et al., 2016). These features provide the perfect opportunity to assess the role of variable resource availability in non-senescent species. Specifically, we focus on the impacts of variable resource availability on planaria population size and population structure, with the rationale for said focus detailed below.

Variation in vital rates (*e.g.*, age-specific survival) dictate population level processes such as population growth rate and extinction risk (Hernández-Yáñez et al., 2022; Lande & Orzack, 1988). Where survival does not decline with age, an indication of negligible or negative senescence, population level processes may differ from populations where survival does decline with age (actuarial senescence). For example, in organisms where survival declines with age, the growth rate of their populations is typically reduced compared to populations of organisms with negligible senescence (Edelfeldt et al., 2019). Crucially, variation in survival trajectories that are caused by age effects may also be driven by environmental factors (Edelfeldt et al., 2019; Roach et al., 2009). Therefore, changes at the population level (*e.g.*, size) due to variable resources can be an indicator of changes in survival trajectories of individuals and potentially “senescence” status. Population-level assessments of the impact of variable resources on survival trajectories are a well-established approach in organisms such as yeast (*Saccharomyces cerevisiae*, Jiang et al., 2002). In yeast, lifespan is assessed by the number of times cells within a multicellular colony produces daughter cells through budding (Jiang et al., 2002). Such an approach is then also amenable to assess the impacts of variable resources on mean life expectancy of a population of other species whereby clonal reproduction is a key life history trait (*e.g.*, planaria, Malinowski et al., 2017). Furthermore, focusing on the population level provides the opportunity to assess how variable resources shape population structure. Indeed, population structure can play a key role in the dynamics of a population through the changes of the ratio of large/old individuals to smaller/younger individuals (Benton & Grant, 1999; Caswell, 2010).

Here, we develop a full factorial design where we experimentally manipulate three different dimensions of resource availability (quantity, quality, and feeding interval) to examine how resource variability impacts the population structure, population performance, and life history trait trade-offs of two potentially immortal species, the planarians *S. mediterranea* and *D. tahitiensis*. We test three hypotheses regarding the impacts of these three dimensions of resource availability on population size and structure, and underlying life history trade-offs, as detailed in Table 1.

**Table 1.**
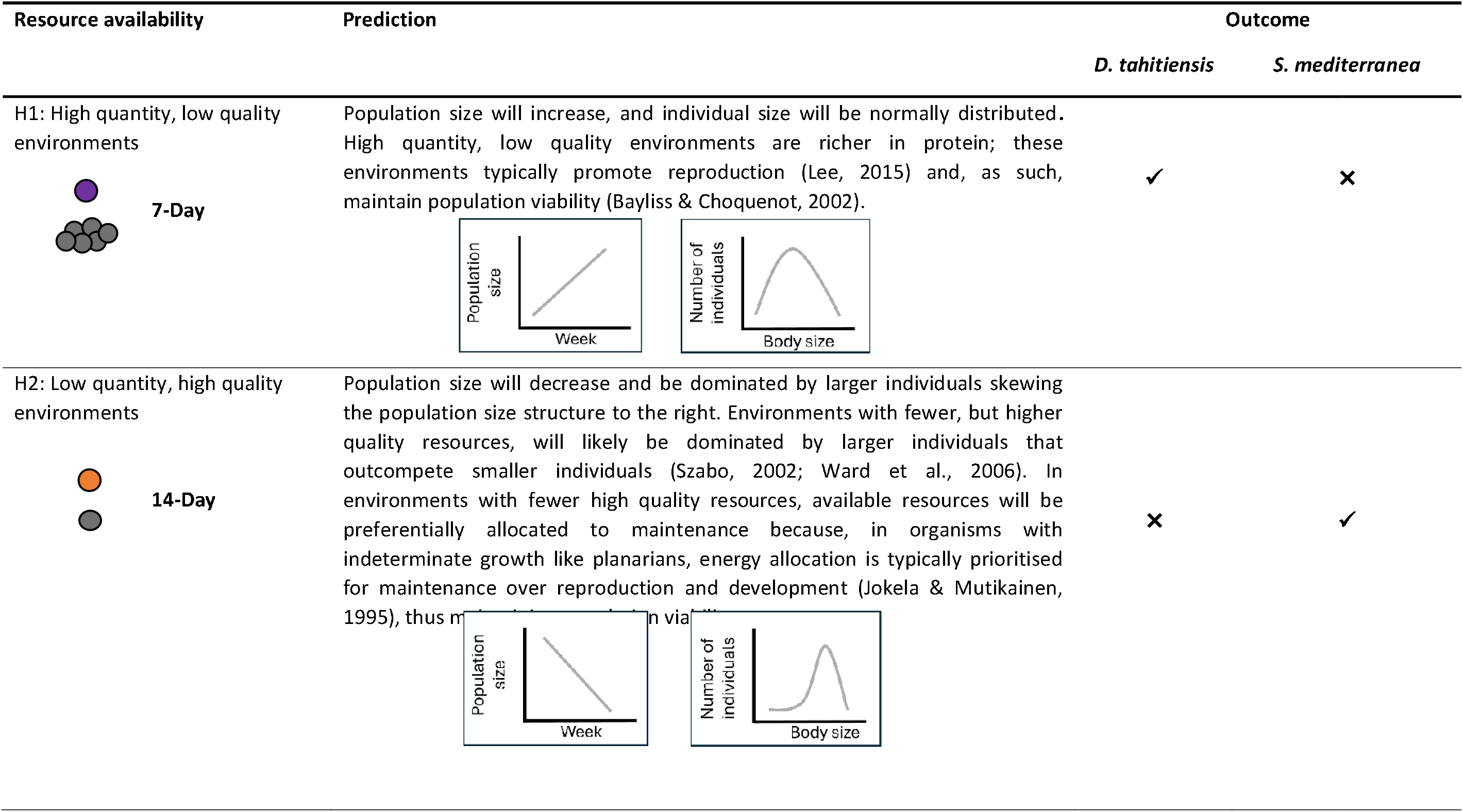
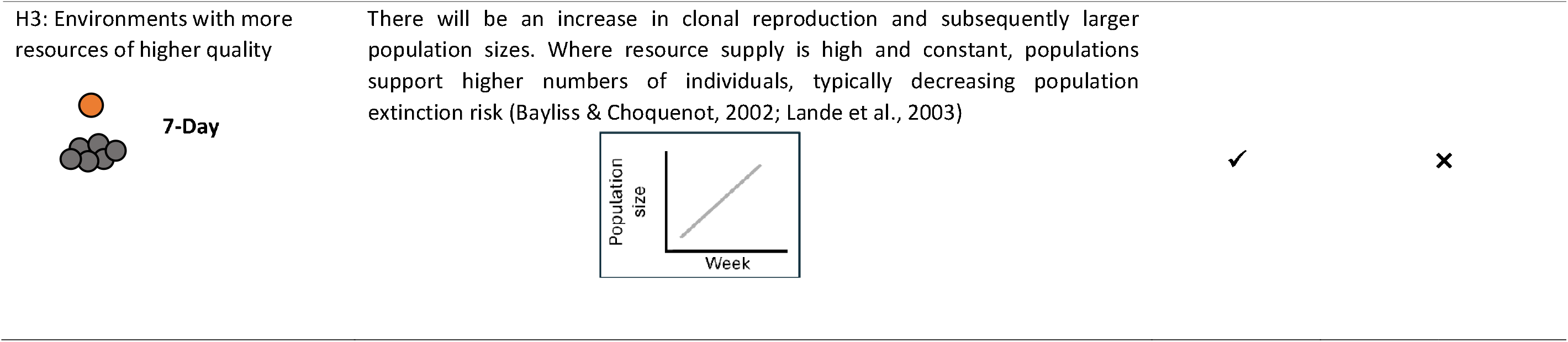
Hypotheses of the impact of the variability in three different dimensions of resource availability (quantity, quality, and feeding interval) on population structure, population performance, and life history trait trade-offs in two species of Planaria; *Schmidtea mediterranea* and *Dugesia tahitiensis*. Here we show results for each species, supporting (✓) or rejecting (✕) each hypothesis. High quality (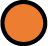); low quality (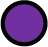); high intake (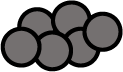); reduced intake (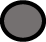) ; feeding intervals every 14 days (14-Day); feeding intervals every seven days (7-Day).

## METHODS

### Study system

To examine how resource variability impacts the population structure, population viability, and life history trait trade-offs, we chose planaria as a study system due to their key life history traits. Notably, the majority of planarians are capable of restorative and physiological regeneration due to injury or damaged/sick cells (Elliott & Sánchez Alvarado, 2013). Their ability to replace damaged or sick cells contributes to the typically long-lived life history of planarians (Elliott & Sánchez Alvarado, 2013). Indeed, under favourable, constant conditions, planaria are theoretically immortal (Elliott & Sánchez Alvarado, 2013). Planaria are carnivorous and feed on live arthropods, annelids, and molluscs. However, planaria also scavenge on individuals that have recently died (Abel et al., 2022; Vila-Farré & C. Rink, 2018). Prey preference has also been shown in several planaria species (Vila-Farré & C. Rink, 2018). Nonetheless, this preference can vary among localities and when the preferred prey is scarce (Vila-Farré & C. Rink, 2018). Importantly, individual planarians are also able to shrink when faced with unfavourable conditions (González-Estévez, 2009).

Here, we study responses to variability in resource availability by *Schmidtea mediterranea* and *Dugesia tahitiensis*, two closely related, freshwater planarian species from the order Tricladida. Both species are free-living and naturally occur in slow running and still water, but are easily reared in the lab (Sousa & Adell, 2018). The two species also differ in size, with *D. tahitiensis* being larger (≈10mm in length) than *S. mediterranea* (≈5mm). *S. mediterranea* can reproduce sexually and asexually, whereas *D. tahitiensis* is an obligate asexual. We used asexual lines of *S. mediterranea* for our experiments to directly compare the two species. Asexual reproduction in planaria occurs via binary fission, which results in two genetically identical clones of smaller size from the fissioning individual (Malinowski et al., 2017). Details of stock culture maintenance are described in the Supporting Information.

### Experiment

To test our hypotheses regarding how variability in resource availability impacts populations of *S. mediterranea* and *D. tahitiensis* (Table 1), we randomly assigned populations to a full-factorial design for each of the two species. This design included three resource factors: quality, quantity, and feeding interval (Fig. 1, Box 1). Quality of resource had two levels, differing in energy content and relative protein-to-carbohydrate ratio: high quality (high carbohydrate level, HQ hereafter) vs. low quality (low carbohydrate level, LQ hereafter). The high quality diet was organic calf’s liver and the low quality diet bloodworm. The protein-to-carbohydrate ratio indicates resource quality, with low carbohydrate-to-protein content corresponding to poorer quality (Lee et al. 2008). The two diets differed in their energy content (109 kcal/100g in the liver vs. 14 kcal/100g in the bloodworm) and in their relative protein-to-carbohydrate ratios (1:0.35 in liver vs. 1:0.17 in bloodworm). Quantity of resources had three levels, which differed in the amount of calories available in each feeding: standard calorie intake (*SI*: 0.001g/individual, representative of intake to maintain laboratory populations (Jochen, 2018)), restricted calorie intake (*RI*: 0.0005g/ind.), hyper-calorie intake (*HI: ad libitum* which was at least 10 fold, approx. 0.01g/individual, that of the SI). Different combinations of resource quantity and quality were provided at two different intervals: once every seven days (*7D*), or every 14 days (14D). The total amount of resources provided over a 14-day period was equivalent for both feeding interval treatments (i.e., one 14-day interval equalled two 7-day intervals) (see Fig. 1). As such, across the most extreme diet treatments, there is over a 300-fold difference in calories provided (per individual per unit time). Within our experimental setup, the equivalent to a control would be either the high quality (*HQ*), standard intake (*SI*), seven days interval (*7D*) treatment (*i.e., HQ-SI-7D*) or the low quality (*LQ*), standard intake (*SI*), 14 days interval (*14D*) treatment (i.e., *LQ-SI-14D*). Here, we determine resources provided at an interval of every seven days (*7D*) of a high quality non-restricted (standard intake for the organism) diet as our control (i.e., *HQ-SI-7D*).

**Figure 1.**
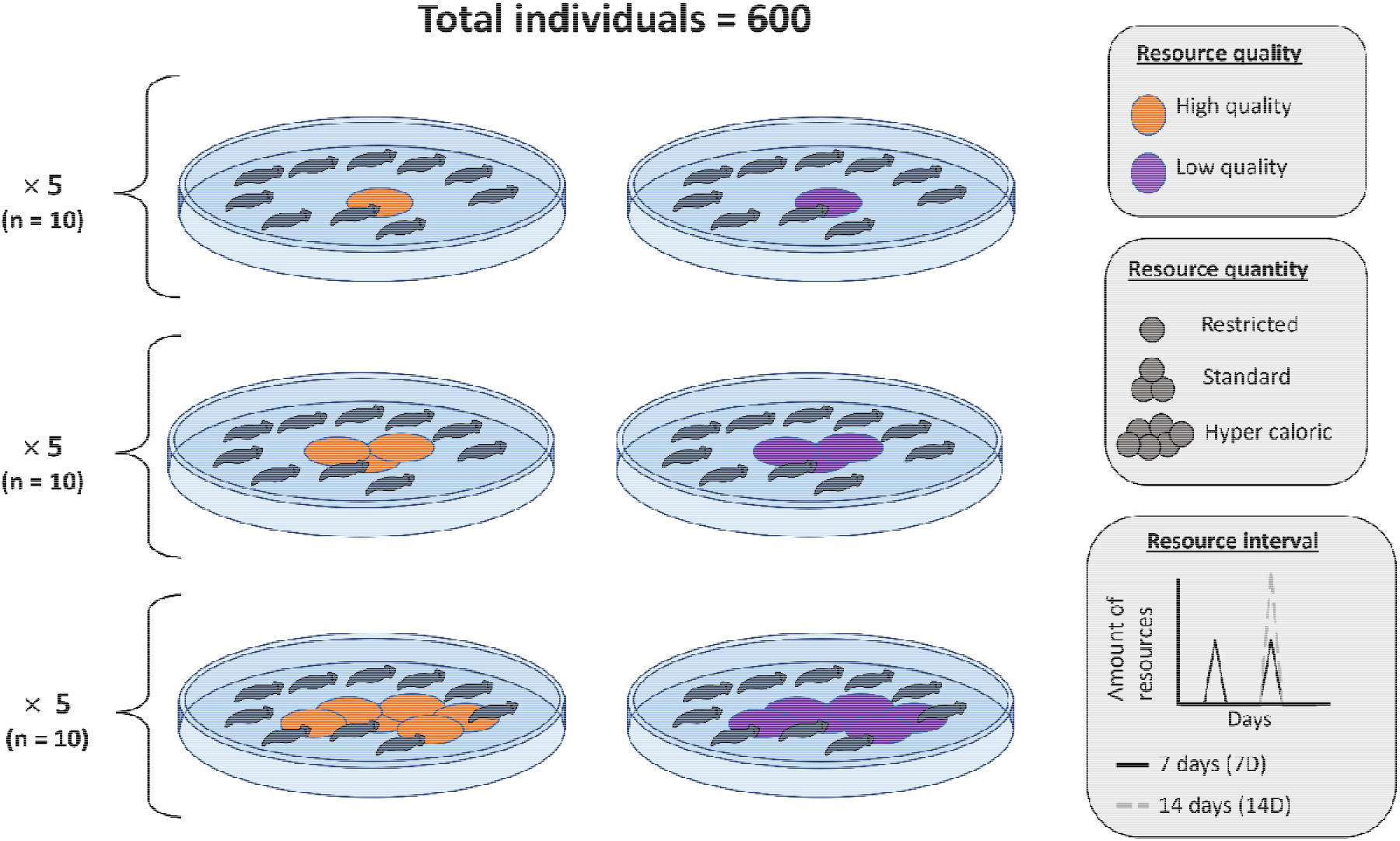
Full factorial experimental design to examine how three key dimensions of resource (quality, quantity, and interval of resource availability) shape population size, population structure and life history trait trade-offs of two flatworm species, *Schmidtea mediterranea* and *Dugesia tahitiensis*. Resource quality has two levels (top right panel): high quality (HQ; orange dots) and low quality (*LQ*; purple dots). Resource quantity has three levels (middle right panel): restricted intake (*RI*; single grey dot), standard intake (SI; three grey dots) and hyper caloric intake (HI; six grey dots). Resource interval has two levels (bottom right panel): every seven days (*7D*; black line), every 14 days (*14D*; grey line). Each population was initiated with 10 individuals and replicated five times resulting in a total of 600 individuals and 60 populations per species. Populations were counted and photographed weekly.

To evaluate interactive effects of resource availability on population structure, population viability, and life history trait trade-offs, we replicated all treatments in the full factorial (n = 12 levels: 2 qualities: HQ vs. LQ; 3 quantities: *RI, SI & HI*; 2 intervals: *7D vs. 14D*) five times for each species. A replicate consisted of a petri dish (25mm deep × 150mm Ø) containing 10 clonal individuals each, henceforth referred to as a population (total n = 600 ind./spp.). For each population, individuals were randomly isolated from one of the three stock cultures and placed in the population 7-8 days before the start of the experiment. Importantly, as individuals of both species are clonal, and stock cultures were initiated with individuals from the same cohort, individuals within each population were of the same chronological age. During this time, individuals were not fed to ensure individuals across all treatments were in a similar starved state at the start of the experiment. Treatment populations were randomised into three blocks containing 20 populations per species per block. For each species, all experimental blocks lasted 19 weeks (133 days). Blocks were staggered in three different start dates due to the logistical constraint of sampling all 60 populations on the same day, as such staggered blocks allowed a more logistically feasible sampling of 20 populations per day. Block 1 of the experiment started on the 4^th^ February 2020, and block 2 on the 30^th^ April 2020. In block 3, the start date for *S. mediterranea* and *D. tahitiensis* differed due to low numbers available for *S. mediterranea* from the stock populations. As such, block 3 for *D. tahitiensis* started on the 1 st May 2020 and for *S. mediterranea* on the 19 th June 2020. Part way through the experiment, due to the COVID-19 pandemic, the location of the experiment was changed within 2 hours, while retaining the same environmental conditions, see Fig S1 in the Supporting Information.

Within each population, every week, population extinction was determined, and asexual reproduction (clonal reproduction) estimated by counting all living and dead individuals. The weekly determination of population extinction and estimation of reproduction was done to determine changes in population size. The expectation is that, in H1 and H3 (Table 1), risk of population extinction will decrease due to increased reproduction, while in H2 (Table 1) extinction risk will decrease due to investment in maintenance. Additionally, size (area in mm^2^) of each individual flatworm within each population was measured weekly to determine growth/shrinkage. Size measurements were calculated from photographs taken of each population (details of image digitization can be found in the Supporting Information). Our expectations here are that for H1, individual size will be normally distributed as the average resource intake among individuals will not be impacted due to the high quantity of resources available. For H2, environments will be dominated by larger individuals due to competitive effects; and for H3, reduced feeding interval of resources will result in greater rate of size oscillation as more frequent resources does not restrict fissioning events to larger individuals (Nentwig & Schauble, 1974).

### Size distribution analysis

To test impact of variation in resource availability on individual body size, we used the size spectrum of populations as a method of quantifying the distribution of body size (Edwards et al., 2017). Where populations are increasing in size, with increases in small individuals, this implies increased reproductive output. To do so, we calculated the size spectra slope values (b) for each population, of both species, across treatments following Carvalho et al. (2021). Briefly, b describes the relative abundance of a population in relation to body sizes, thus providing a good proxy to quantify the distribution of body size within a population. A steeper (more negative) b indicates fewer large-bodied and/or more small-bodied individuals, which is an indication of a right-skewed size population structure. Values of b can vary substantially with estimates ranging from -6.75 to -0.40 been reported (Arranz et al., 2021; Carvalho et al., 2021). To estimate *b*, for each species separately, we used the abundance density function (Carvalho et al., 2021):

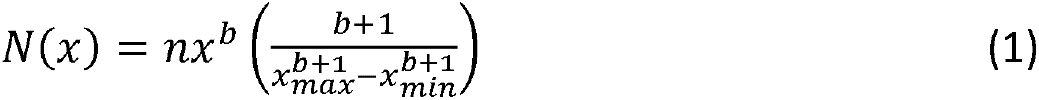

where *n* is the number of individuals, *x* the area of each individual *x_max_* and *x_min_* are the maximum and minimum area of individuals that were digitized. To calculate the maximum and minimum values, all populations for each treatment were pooled and maximum and minimum values calculated separately for each treatment for each week.

For each week of the experiment, *b* values were calculated using the area measurements of each individual within a given population. The *b* value for each of the five populations of each treatment was calculated. As such, here *b* relates to how the populations responds to the varying quality, quantity, and feeding interval of resources provided. Next, we applied a Bayesian framework to determine changes in the mean *b* values using the “bmrs” package (Bürkner, 2017) in R version 4.0.3 (R Core Team, 2020). Given that the estimated b values are based on size values, there is the potential that the b values may show a right-skewed distribution due to the ability of planaria to shrink. Thus, we initially compared models with a Gamma distribution to that of a Gaussian distribution to identify the model with the best fit. We identified that in the case of both *S. mediterranea* and *D. tahitiensis* a model with a gaussian distribution had the best fit (Supporting Information, Table S2). The final model used to model the response variable *b*, had a Gaussian distribution and adaptive priors with treatment (categorical with 12 levels), week (19 weeks) and block (3 blocks) as fixed variables and population (60 populations) as a random effect. Presented are the median estimate and 50% and 95% compatibility intervals of the posterior probability. Compatibility intervals can be considered equivalent to confidence intervals in a frequentist approach (Gelman & Greenland, 2019). See Supporting Information for model details and Table S3 & S4 for final priors of each model.

### Population size analysis

To determine the impact of variation in resource availability on population size, we calculated changes in weekly population size for each species. We applied a Bayesian framework to determine changes in population size using the same “bmrs” package as before. Population size was modelled with a Poisson and Gaussian distribution to determine best fit. In the case of *S. mediterranea* the model with the best fit was identified as one with a Gaussian distribution (Supporting Information, Table S5) and adaptive priors, with treatment (categorical with 12 levels), week (19 weeks) and block (3 blocks) as fixed variables, population as a random effect (60 populations) and population size the response variable. In the case of *D. tahitiensis*, the model with the best fit had a Poisson distribution with a log link function (Supporting Information, Table S5) using adaptive priors, with treatment (categorical with 12 levels), week (19 weeks) and block (3 blocks) as fixed variables, population as a random effect (60 populations) and population size the response variable. As before, median estimated values are presented with 50% and 95% compatibility intervals. See Supporting Information for model details and Table S6 & S7 for final priors of each model.

Crucially by analysing changes in population size and population structure (as identified by the size distribution analysis), we can provide insights in potential reproductive output given our knowledge of the biology and life cycle of both species (Carter et al., 2015; González-Estévez, 2009; Malinowski et al., 2017; Nentwig & Schauble, 1974). For example, a population that is increasing in its size with increases in the number of small individuals implies increased reproductive output. In a population where individuals are shrinking, fissioning (reproduction) is not occurring and populations are not increasing. As such there will not be overlap with the size distributions of small individuals due to reproductive output and small individuals as a result of shrinking.

## RESULTS

### Body size distributions

Over time, size spectra slope values (*b*) for *S. mediterranea* became more negative for most treatments (Fig. 2), implying a more right-skewed population. The consistent decline in b values indicates a change in the size structure of the populations with an increase in smaller individuals and fewer larger individuals. Overall, 14-day interval treatments had more negative *b* values than 7-day interval treatments (Fig. S3). In the high-quality treatments, when compared to *HQ-SI-7D* (our control), only the 14-day feeding interval high intake (*HQ-HI-14D*) and 14-day feeding interval reduced intake (*HQ-RI-14D*) treatment had smaller b values and as such more right-skewed size structure, with the *HQ-RI-14D* treatment fulfilling the size prediction of H2 (Table 1). This outcome suggests that in high quality treatments populations changed in size structure with an increase in the number of small individuals and a decrease in the number of larger individuals over time (Fig. 2). When changes in structure were found, this change in structure was more prominent in treatments where feeding interval was 14 days. However, in the case of the low-quality treatments, only the 7-day feeding interval reduced intake (*LQ-RI-7D*) treatment did not show a change in size structure that differed from that of *HQ-SI-7D* (Fig. S3). All other low quality treatments showed a greater change in size structure (i.e., more negative b values) with more right-skewed populations over time (Fig. 2). In the case of *LQ-HI-7D*, the size prediction of H1 was not fulfilled as size was predicted to be normally distributed.

**Figure 2.**
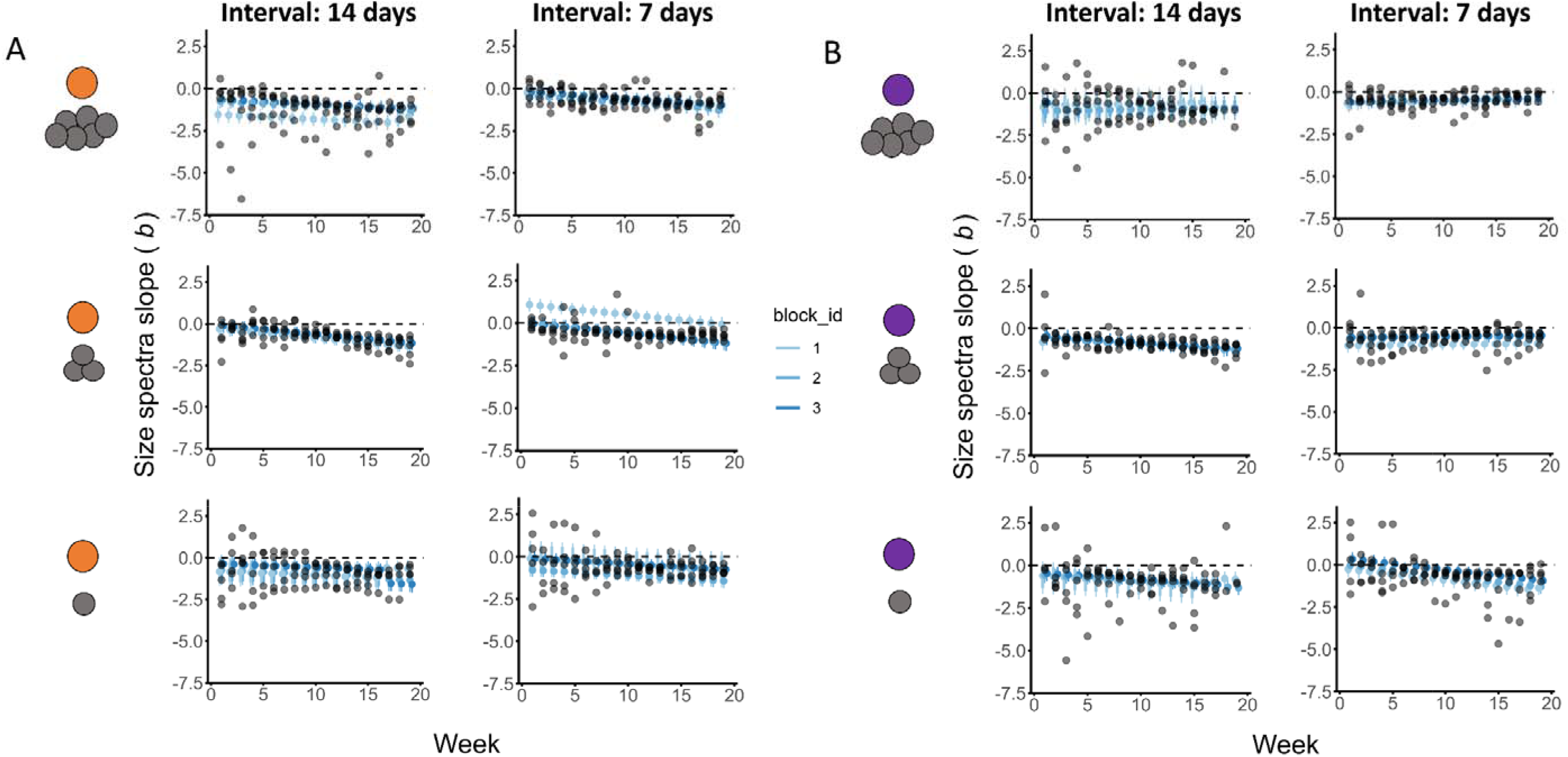
Population structure in *Schmidtea mediterranea* populations changes across time in most treatments. Weekly size spectra slope values (*b*) for *S. mediterranea*. A) Calculated size spectra slope (*b*) values (black points) for all populations for high quality (HQ, 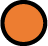) treatments over 19 weeks. B) Calculated size spectra slope (*b*) values (black points) for all populations for low quality (*LQ*, 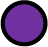) treatments over 19 weeks. Blue points indicate estimated *b* values with compatibility intervals (0.95) for each block (block 1 – light blue; block 2 – blue; block 3 – dark blue). Left panels in each graph indicate feeding intervals every 14 days and right panels feeding intervals every seven days. Top panel represent high intake (*HI*, 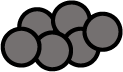), middles panel standard intake (*SI*, 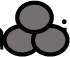) and bottom panel reduced intake (*RI*, 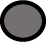) treatments.

For *D. tahitiensis*, size spectra slope values (b) varied less in the 7-day feeding interval treatments than the 14-day feeding interval treatments (Fig. 3). When compared to *S. mediterranea*, *D. tahitiensis* populations were less right-skewed. When assessing the outcomes in terms of the size predictions of H1 and H2 for *D. tahitiensis*, treatment *LQ-HI-7D* fulfilled the predictions of H1 as body size was normally distributed (Table 1). However, for treatment *HQ-RI-14D* the size predictions of H2, a right-skewed population, was not fulfilled (Table 1). In both species experimental block 1 differed from block 2 and block 3 in several treatments (Fig. S3 & S4). However, this pattern was not consistent between treatments and species (see Fig. 2 and 3, light blue points). Still, even with the block differences, the trends of all blocks were the same (increasing, decreasing, or stable). Overall, there was no consistent trend in how b values across treatments differed to that of *HQ-SI-7D*. As such, treatment effect does not predict variation in size structure over time.

**Figure 3.**
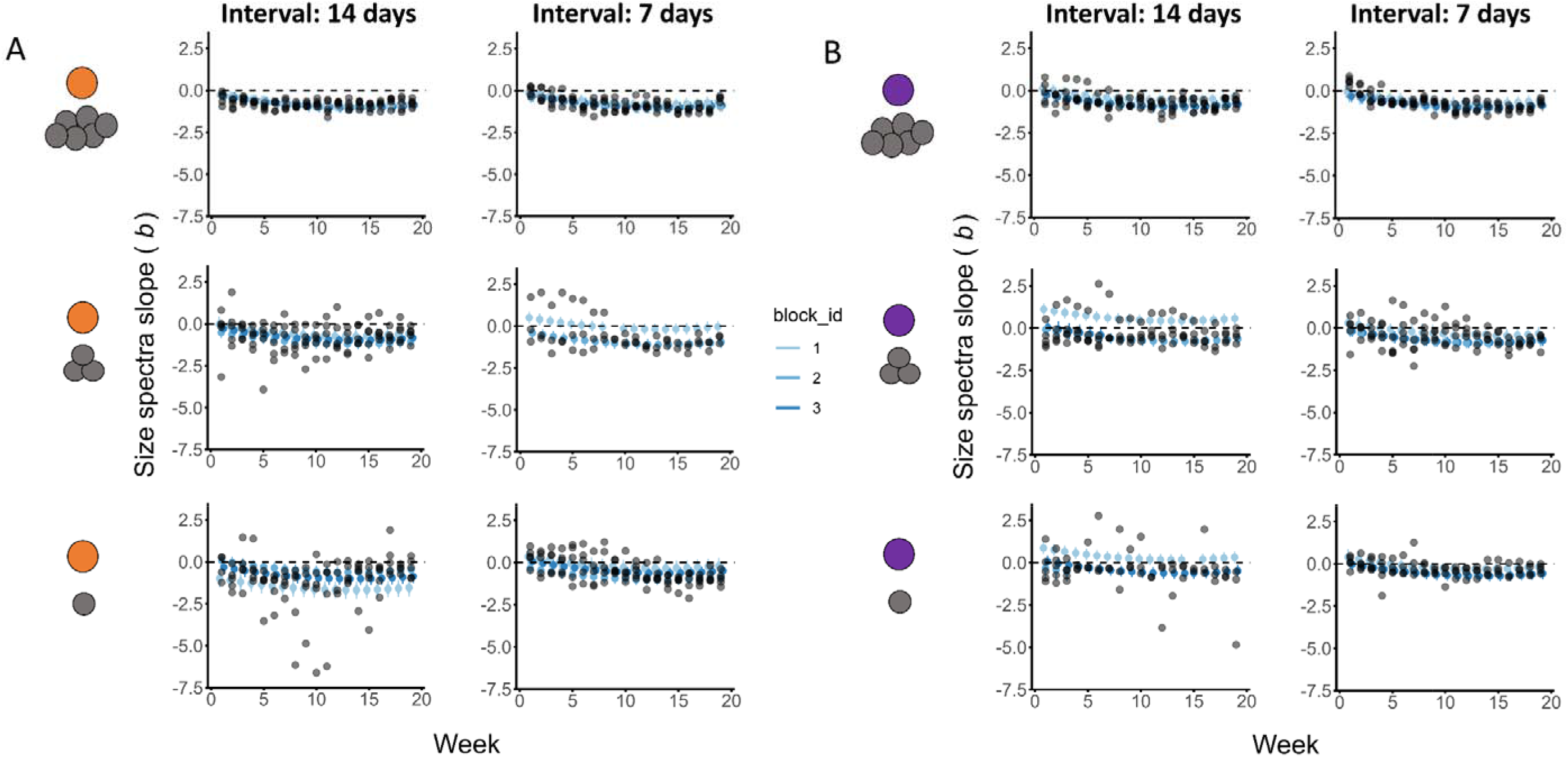
Population structure in *Dugesia tahitiensis* populations has little change in structure across calorie restriction treatments. Weekly size spectra slope values (b) for *D. tahitiensis*. A) Calculated size spectra slope (b) values (black points) for all populations for high quality (*HQ*, 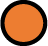) treatments over 19 weeks. B) Calculated size spectra slope (b) values (black points) for all populations for low quality (*LQ*, 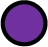) treatments over 19 weeks. Blue points indicate estimated b values with compatibility intervals (0.95) for each block (block 1 – light blue; block 2 – blue; block 3 – dark blue). Left panels in each graph indicate feeding intervals every 14 days and right feeding intervals every seven days. Top panel represent high intake (*HI*, 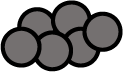), middles panel standard intake (*SI*, 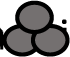) and bottom panel reduced intake (*RI*, 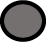) treatments.

### Population trends

Overall, *D. tahitiensis* populations had higher numbers than populations of *S. mediterranea* at the end of the experiment (Fig. 4 & 5). Population size of *S. mediterranea* varied between treatments that differed in resource quantity (HI: high intake, SI: standard intake, RI: reduced intake) and interval of feeding (7D: every seven days, 14D: every 14 days) but was consistent across resource quality (*HQ*: high quality, *LQ*: low quality) (Fig. 4A & B). The exception being the high intake resource quantity treatments with a 7-day feeding interval (HI-7D). In this case, the *HQ-HI-7D* treatment, where the resource was of a high quality, the population declined compared to the *LQ-HI-7D* treatment, where the population increased over time (Fig. 4A & B, Fig. S5). All populations where resource was provided every 14 days showed a decrease in population size over time compared to populations when feeding interval was every seven days, where counts either stayed constant (*HQ-RI-7D*) or increased over time (*HQ-SI-7D, *LQ-HI-7D*, LQ-SI-7D*) (Fig. 4A & B Fig. S5). The exception here was the *HQ-HI-7D* treatment, where population size declined over time. The decline in *HQ-HI-7D* does not fulfil the prediction of H3 where the expectation was an increase in reproduction and a larger population size. However, the decrease in population size of treatment *HQ-RI-14D* supports the population size prediction of H2. As such, the outcome of *HQ-RI-14D* fulfils the full prediction of H2 that population size will decrease and body size will be right-skewed.

**Figure 4.**
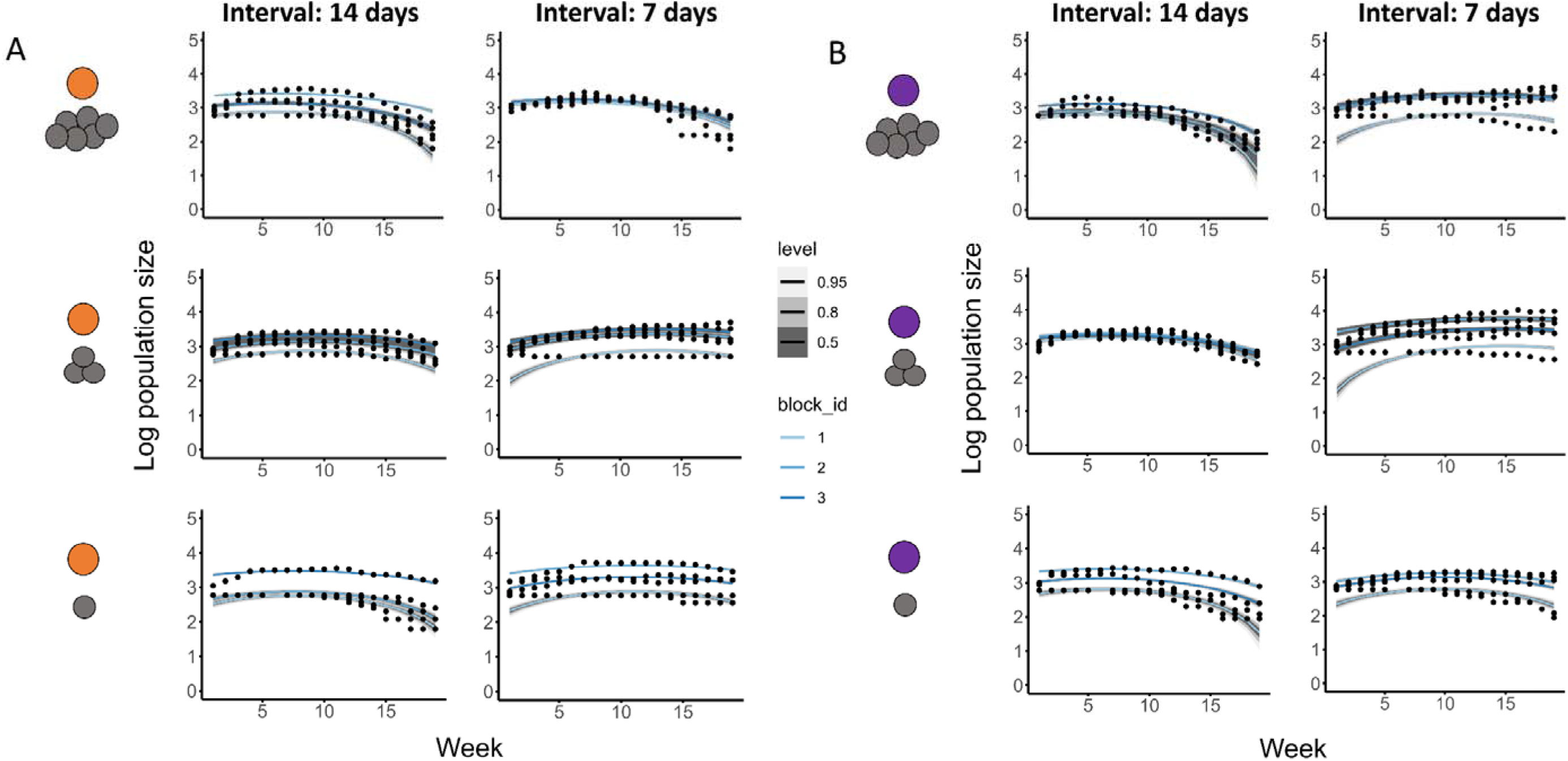
Population size for *Schmidtea mediterranea* varies but with no clear patterns across calorie restriction treatments. Weekly population size of *S. mediterranea* over the 19 weeks for A) high quality (*HQ*, 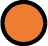) treatments and B) low quality (*LQ*, 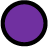) treatments. Black points indicate raw data of weekly population size; blue lines indicate estimated population size for each block (block 1 – light blue; block 2 – blue; block 3 – dark blue). Shaded grey regions indicate compatibility intervals: light grey 0.95, grey 0.8, dark grey 0.5. Left panels indicate feeding intervals every 14 days and right panels feeding intervals every seven days. Top panel represent high intake (*HI*, 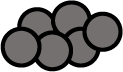), middles panel standard intake (*SI*, 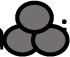) and bottom panel reduced intake (*RI*, 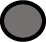) treatments.

In *D. tahitiensis*, populations either increased or remained stable across all treatments (Fig. 5A & B). When comparing HQ and LQ treatments, HQ treatments with feeding intervals of seven days, regardless of resource quantity, had a higher population size than LQ treatments with 7-day feeding intervals, except for *HQ-HI-7D* and *LQ-HI-7D*, which had similar counts. When resource was provided every 14 days, there was no clear trend; the *HQ-HI-14D* treatment had lower counts than the LQ-HI-14D treatment, *HQ-SI-14D* had higher counts than *LQ-SI-14D* and *HQ-RI-14D* had similar counts to LQ-RI-14D (Fig. 5A & B, Fig. S6). In the case of *HQ-RI-14D* population size increased which did not fulfil the population predictions of H2. For treatments *LQ-HI-7D* and *HQ-HI-7D* population size increased, thus fulfilling the population predictions of H1 and H3 respectively. For both species, experimental block 1 (light blue lines, Fig. 4 & 5) had population growth rates lower than block 2 and block 3. However, for all treatments, the population response (increasing, decreasing, or stable counts) of block 1 did not differ from that of block 2 or block 3 (Fig. 4 & Fig. 5, Fig. S5 & S6).

**Figure 5.**
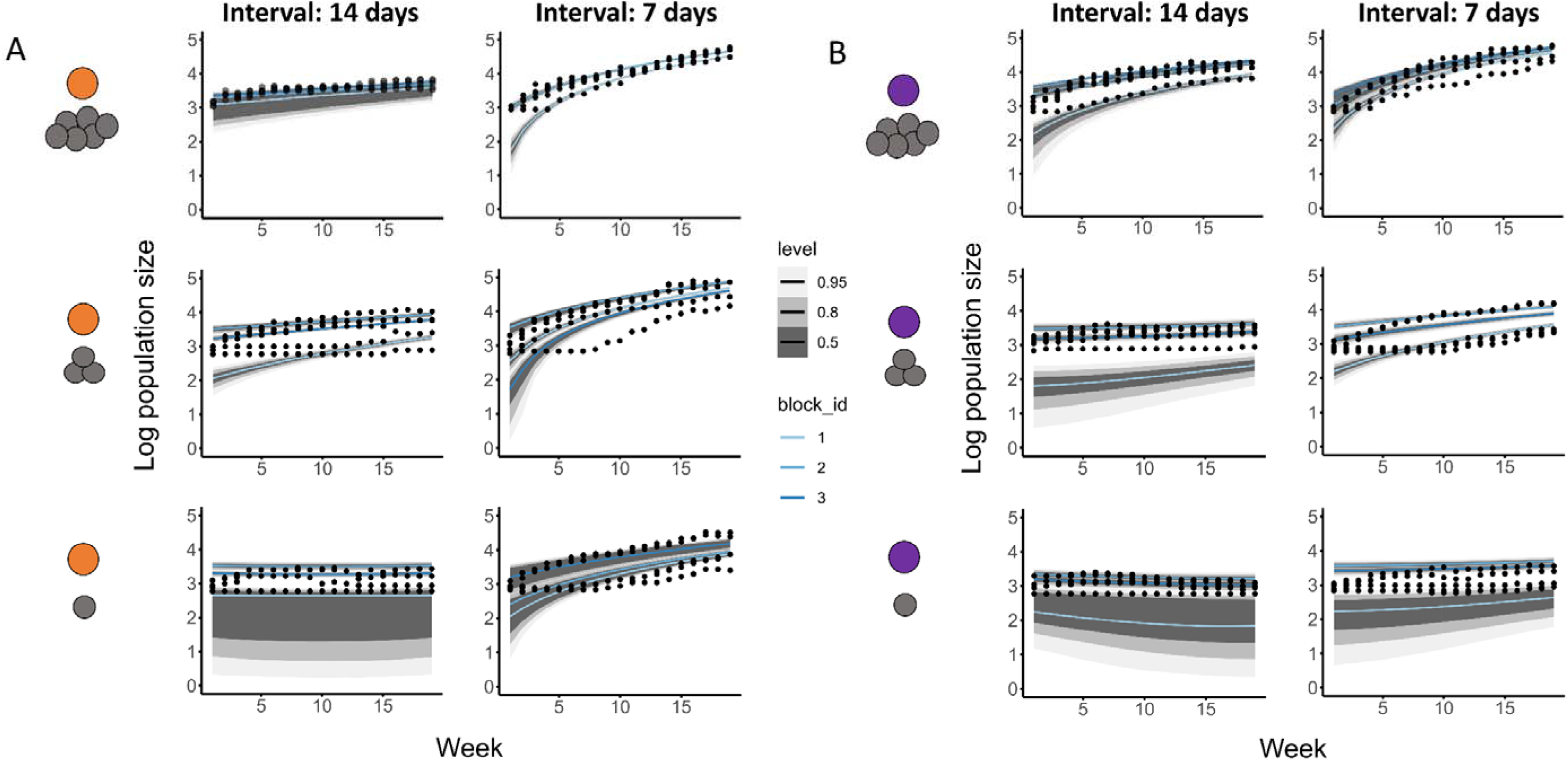
Population size for *Dugesia tahitiensis* varies but with no clear patterns across calorie restriction treatments. Weekly population size of *D. tahitiensis* over the 19 weeks for A) high quality (*HQ*, 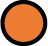) treatments and B) low quality (*LQ*, 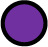) treatments. Black points indicate raw data of weekly population size; blue lines indicate estimated population size for each block (block 1 – light blue; block 2 – blue; block 3 – dark blue). Shaded grey regions indicate compatibility intervals: light grey 0.95, grey 0.8, dark grey 0.5. Left panels indicate feeding intervals every 14 days and right panels feeding intervals every seven days. Top panel represent high intake (*HI*, 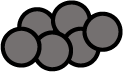), middles panel standard intake (*SI*, 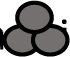) and bottom panel reduced intake (*RI*, 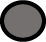) treatments.

## DISCUSSION

Compared to senescent species (Lemanski et al., 2020; Pardo et al., 2013), a comprehensive understanding of the impact of variable resource availability on species that show negligible or negative senescence is lacking (but see Schaible et al., 2011). Here, we contribute to this gap of knowledge by incorporating variation in resource availability in two non-senescent species: the flatworms *S. mediterranea* and *D. tahitiensis* (Sahu et al., 2017). Our findings show how, when multiple factors of resource availability interact (i.e., quality, quantity and frequency), these non-senescent species show population-level responses expected from suboptimal conditions of stress. Specifically, in *S. mediterranea*, we observe suboptimal conditions of stress through population decline and individual shrinkage when resources are made available are every 14 days, regardless of the resource quality or quantity. However, conditions of stress in *Dugesia tahitiensis* are less frequent and occur only when resources are reduced, regardless of resource intervals, with only individual-level shrinkage being noted as evidence of stress, since we observe no population decline.

The way these factors of resource availability impacts individuals within populations is context dependent in our experiment. The high population viability (only two of the 20 populations went extinct in our experiment, both were *S. mediterranea*) and variable response in reproduction (as proxied by population size and structure) across treatments is also likely driven by the specific life history strategy of our species. Generally, planarians can restore their own damaged cells (Elliott & Sánchez Alvarado, 2013). However, restoring damaged cells is costly and requires resources (Akashi & Gojobori, 2002). Naturally, if most resources are used for maintenance, fewer resources will be available for reproduction. The high population viability we report here in response to the variable resource availability is indicative of the fundamental trade-off between survival and reproduction under variable environmental conditions (Stearns, 1992). Here, theory predicts that, in long lived species such as our planarians (Elliott & Sánchez Alvarado, 2013), resources will be primarily invested in maintenance of individuals and not reproduction (Erikstad et al., 1998; Sæther et al., 2013). As highlighted earlier, reproduction in planaria occurs via binary fission, and so individuals in our experimental setup are clonal and thus of the same age. As such, any changes in the structure and size of a population can be thought of as a reflection of the survival-reproduction trade-off. A less right-skewed population, where there are fewer smaller individuals, is indicative of reduced reproduction as new recruits are smaller. In these populations of reduced reproduction, individuals invest in maintenance that results in reduced/no growth with little change in body size. In general, for *S. mediterranea*, such a survival-reproduction trade-off can be observed across the 7-day feeding treatment, but not when the feeding interval was every 14 days, where population size decreased, indicating reduced reproduction and reduced survival of individuals. For *D. tahitiensis*, only populations of low quantity and with a feeding interval every 14 days show a survival-reproduction trade-off of high survival and reduced reproduction. Such context-dependant trade-offs have also been shown in Hydra magnipapillata, another non-senescent species (Schaible et al., 2011). There, the authors found a clear maintenance-reproduction trade-off when resources and temperature were constant, whereby investment into reproduction resulted in reduced survival. However, under fluctuating environmental conditions, the authors reported no clear maintenance-reproduction trade-off. This context-dependent survival-reproduction trade-off then begs the question, where there is an apparent lack of trade-off between survival and reproduction (see Metcalf 2016), do we see a trade-off with body size?

The size structure of our populations varied across treatments, but not in a way that was consistent with our hypotheses (Table 1). For *S. mediterranea*, populations become more right-skewed across treatments, with two exceptions in the 7-day feeding interval treatment. The right-skewed populations describe an increase in the number of smaller individuals and decline of larger individuals in the populations (Carvalho et al., 2021). Such changes in population structure can occur when there is increased reproduction, as more recruits of a smaller size enter the population. *S. mediterranea* reproduce asexually through fission, where larger individuals fission into two, with the posterior product (new recruits) generally smaller than the anterior fission product (now a smaller individual too) (Peter et al., 2001). Thus, fissioning results in fewer larger individuals and an increase in the number of smaller individuals. However, the right-skewed size structure due to reproductive events will only be reflected in populations that are increasing in number as in these populations no mortality of individuals was observed. Where *S. mediterranea* populations show a decline or remain stable in size, the change in structure is not driven by decreased reproduction. Instead, the life history of planaria seems to be driving the response. A key trait in the life history of planaria is the ability to shrink when resources become scarce (Mangel et al., 2016; Rink, 2013).

The ability to shrink allows individuals of some species to survive resource-scarce periods of time until conditions improve (Ebert, 1967; Marinovic & Mangel, 1999; Wikelski & Thom, 2000). In planaria, when individual shrinkage occurs, cell numbers reduce through autophagy during which energy resources are supplied to stem/regenerative (neoblasts) cells (González-Estévez, 2009; Romero & Baguñà, 1991). This process ensures that maintenance levels can be met and as individuals attempt to maintain homeostasis, fissioning (asexual reproduction) does not occur. Furthermore, in *S. mediterranea* and in sister species of *D. tahitiensis* (i.e., D. japonica and D. dorotocephala) the probability of fissioning declines as individuals get smaller, with a minimum size where fissioning can occur (Carter et al., 2015). In the case of sexually reproducing planaria, reproductive organs are often consumed during shrinkage (Saló, 2006), thus preventing further reproduction. Shrinkage is a strategy found in several taxa and been suggested to mitigate the impact of fluctuating or variable environments on individual survival (Ebert, 1967; Marinovic & Mangel, 1999; Salguero-Gómez & Casper, 2010, 2011). Indeed, shrinkage maintains relatively high levels of survival during periods of resource scarcity compared to strategies where shrinkage does not or cannot physically occur. As such, the high levels of survival ensure population viability. For instance, in the Galápagos marine iguanas (*Amblyrhynchus cristatus*), individuals shrink up to 20% of their body length when nutritional resources are low following El Niño (Wikelski & Thom, 2000); those individuals that shrink more have a higher rate of survival, suggesting the adaptive value of shrinkage. When nutritional resources improve during La Niña events, body length then increases again, as is the case with planarians when they receive increased nutritional resources after periods of low nutrient availability (González-Estévez, 2009; Thommen et al., 2019).

The impact of variable resources on body size due to shrinkage observed in *S. mediterranea* populations is unlikely occurring in populations of *D. tahitiensis*. Fewer *D. tahitiensis* treatments showcased populations becoming more right-skewed (i.e., more smaller individuals) over the course of the experiment. There, where populations become more right-skewed, populations do not show a decline in population size and, unlike the case with *S. mediterranea*, changes in population structure are likely due to fissioning. Populations of *D. tahitiensis* that have a less right-skewed size structure increased in size during the experiment. Within these population, individuals are likely experiencing increased competition for resources. Increased resource competition among individuals results in contest competition, where an individual interferes with other individuals to obtain resources (Nicholson, 1954). In competitive environments, smaller individuals are typically outcompeted by larger individuals within the population as larger individuals tend to be more dominant (Petersson & Järvi, 2000; Stewart & Tabak, 2011). In our study, contest competition results in differences in average resource intake among individuals, with larger planarian individuals able to consume more resources per feeding event. As such, competition between two individuals of planarian naturally results in the larger of the two growing more. The competitive exclusion of smaller individuals thereby increases the number of larger individuals in the population, resulting in less right-skewed populations. Here, competition for resources resulted in mortality of smaller individuals and faster growth in the surviving, larger individuals. Evidence of smaller individuals being outcompeted by larger individuals has been observed in several species, including other planaria (Reynoldson, 1964; Reynoldson & Sefton, 1972). For example, in a 20 month study of a lake population of *Planaria torva*, Reynoldson and Sefton (1972) found changes in the size distribution of individuals in the population that were attributed to intraspecific competition for resources. Similarly, in the brown trout (*Salmo trutta*), Petersson & Järvi (2000) observed that sea-ranch trout, which is 13% larger than its wild-type counterpart, had higher dominance than the latter.

The high population viability reported here highlights the ability of planaria to mediate impacts of variation in resource availability. However, as mentioned before, the mechanisms that allow shrinkage and regeneration of individuals do not preclude individuals from suffering suboptimal conditions of stress and ultimately dying. Said resource-availability driven stress is observed in the few *S. mediterranea* populations that went extinct and those populations declining in numbers. Why such extinction events and declining numbers only occurred in populations of *S. mediterranea* and not *D. tahitiensis* can be attributed to differences in their ability to regenerate. Regeneration is not uncommon within the Metazoa, with almost every phylum having a species with this ability (Alvarado, 2000). Furthermore, regeneration can be classified into two categories highlighting the variability of the phenomenon: 1) morphallaxis, the regeneration in the absence of active cell proliferation (*e.g., Hydra*) and 2) epimorphosis, the regeneration with cell proliferation (*e.g.*, limb and tail regeneration in vertebrates) (Alvarado, 2000; Morgan, 1901). In the case of planaria, both forms of regeneration are utilised but are context dependent; morphallaxis to restore scale and proportion (*e.g.*, when shrinking occurs) and epimorphosis for the formation of a new organ (*e.g.*, due to a fission event) (Pellettieri, 2019).

The lack of local extinction and declining numbers in the examined populations of *D. tahitiensis*, compared to *S. mediterranea*, is likely due to the former species being able to implement epimorphosis and repair damaged cells more efficiently than the latter (Peter et al., 2001). Indeed, *Dugesia tahitiensis* has one of the highest regenerative and fissioninig capacities and the most neoblasts (adult stem cells found in Platyhelminthes and Acoela) within the Tricladida taxon (Baguñá & Romero, 1981; Peter et al., 2001). High neoblast density in the cellular tissue between the body wall and organs is a prerequisite for efficient regeneration after fission and, when compared to *S. mediterranea*, *D. tahitiensis* have higher cell counts (Peter et al., 2001). Moreover, neoblast self-renewal activity can be impacted by resource availability with reduced resource levels resulting in lowered self-renewal activity (Mangel et al., 2016). Reduced neoblast self-renewal activity will then enhance the negative impacts experienced by *S. mediterranea*. The variation in self-renewal activity brought about by variation in resource availability will ultimately impact survival trajectories of individuals. As such, the fittest individuals are more likely to survive to extreme ages that increases the population’s average survival, a pattern that has also been described in other animals (Vaupel & Yashin, 1985). This age-by-environment interaction has also been reported in some long-lived plants, e.g. in Plantago lanceolata (Roach et al., 2009) and Fumana procumbens (Edelfeldt et al., 2019), where variation in survival trajectories are modulated by resource availability.

No species is truly immortal. The changing environment can play a significant part in the levels of stress and individual faces, regardless of its senescence rates. Here, we contribute to the under-represented empirical demonstration of the impact of variation in resource availability in two species that do not senesce. Our findings support previous work on another non-senescent species (Hydra magnipapillata; Schaible et al., 2011), showing that the impacts of fluctuating resources on the trade-off between reproduction and maintenance is context dependent. Importantly, we highlight that two non-senescent species respond differently to variation in resource availability; one species can suffer population extinction and the other population growth, highlighting differential suboptimal conditions of stress under changing environmental conditions. A clearer understanding of how fluctuating environments, particularly in resources availability, impacts species that show the unique trait of negligible or negative senescence will be key to fully understand the role of this life history trait and ultimately whether anything is indeed immortal.

## Supporting information

SOM

## Acknowledgements

This project was funded by an Oxford John Fell Funds grant to RSG. RSG was supported by a NERC Independent Research Fellowship (NE/M018458/1).

## Author Contributions

RSG conceived the idea; JAD, RSG, and AA designed the methodology; JAD and PH collected and digitized data; JAD analysed data; JAD led the writing of the manuscript with assistance from RSG. All authors contributed critically to the drafts and gave final approval for publication.

## Data Availability

The data in this paper are published as supporting information in the open access version deposited in bioRxiv.org (DOI: 10.1101/2023.01.12.523873; Deere et al. 2022) (data are in multiple formats; *e.g.*, .pdf, .csv, .cpproj).

## BOX 1 Glossary

HQ-HI-7D: Populations provided with high quality (HQ), high intake (HI) resources at intervals of every seven days.

HQ-HI-14D: Populations provided with high quality (HQ), high intake (HI) resources at intervals of every 14 days.

HQ-SI-7D: Populations provided with high quality (HQ), standard intake (SI) resources at intervals of every seven days.

HQ-SI-14D: Populations provided with high quality (HQ), standard intake (SI) resources at intervals of every 14 days.

HQ-RI-7D: Populations provided with high quality (HQ), reduced intake (RI) resources at intervals of every seven days.

HQ-RI-14D: Populations provided with high quality (HQ), reduced intake (RI) resources at intervals of every 14 days.

LQ-HI-7D: Populations provided with low quality (LQ), high intake (HI) resources at intervals of every seven days.

LQ-HI-14D: Populations provided with low quality (LQ), high intake (HI) resources at intervals of every 14 days.

LQ-SI-7D: Populations provided with low quality (LQ), standard intake (SI) resources at intervals of every seven days.

LQ-SI-14D: Populations provided with low quality (LQ), standard intake (SI) resources at intervals of every 14 days.

LQ-RI-7D: Populations provided with low quality (LQ), reduced intake (RI) resources at intervals of every seven days.

LQ-RI-14D: Populations provided with low quality (LQ), reduced intake (RI) resources at intervals of every 14 days.

