## Supplementary material for "Non-senescent species are not immortal: stress and decline in two planaria species": SOM

**This file includes:**

Methods

Supplementary Table 1 - 6

Supplementary Figures 1 - 6

**Methods**

*Stock culture*

To form cultures, a clonal line of the wild type, asexual strain of *S. mediterranea* and *D. tahitiensis* were initiated from long-term cultures maintained by the Aboobaker laboratory in the Department of Zoology at the University of Oxford. The *S. mediterranea* cultures originate from the main laboratory stock used worldwide, first collected from Monjuïc, Barcelona, Spain. *D. tahitiensis* originate from the cultures in the Egger laboratory at the University of Innsbruck, Austria. As these populations increase through binary fission all individuals are clonal. A total of three stock cultures per species were started with 10 clonal individuals each, as such all individuals can be considered of the same cohort and are the same age. To explore the impact of three different dimensions of resource availability (quality, quantity, interval; see ‘Experiment’ below), we increased initial numbers of planarians to ensure a large sample size. Once a week, for three weeks, 10 individuals per species were taken from each culture and two transverse amputations per individual were performed (thus forming three individuals) following methods in Sousa and Adell (2018). In doing so, all individuals were of a similar chronological age and the potential age variance was reduced to three weeks, which is significantly less than that of individuals within the main laboratory stock. Once fissioned, during regeneration, polarity of their body axes is maintained. This means that each fissioned piece conserves the anterior-posterior, dorsal-ventral and medial-lateral axes and, as such, subsequent organogenesis (Elliott & Sánchez Alvarado 2013). Individuals were reared in a mixture of distilled water and sea salt (Instant Ocean sea salt, Aquarium Systems) at a concentration of 0.5g/L. Cultures were fed on organic calf’s liver twice a week; the liver was left for two hours in the containers before being removed to ensure full satiation of individuals within the population. After removal, the water-salt mix was replaced and the containers housing the cultures were cleaned to avoid potential infection through debris build-up. This feeding regime frequency was sufficient to ensure continued fissioning of individuals and increasing numbers in the cultures. Stock cultures were housed in six 30cm long × 30cm wide × 15cm high plastic containers (three for *S. mediterranea*, three for *D. tahitiensis*) within a climate room set at 20°C and kept in 0:24hr light:dark regime. Populations were only exposed to low levels of light for short periods of time during feeding and data collection.

*Experimental setup*

To determine the impact of variation in available resources on population size, population structure and life history trait trade-offs, we conducted experiments incorporating three different dimensions of resource availability (quantity: standard intake (SI), restricted intake (RI), high intake (HI); quality: high-quality (HQ), low-quality (LQ); resource interval: seven days (7D), 14 days (14D)) (see main text for details). As part of the experimental protocol, we always fed treatments the day after data collection (population counts and photographs taken) to ensure counts were reflective of the previous feeding event. We based the amount of resource provided a given week on the population counts of the previous day and the relevant treatment (*i.e.*, total number of individuals multiplied by the resource weight for the specific treatment). All resources were weighed to the nearest 0.0001g using an Adventurer Analytical microbalance (Ohaus). After feeding, all the water-salt mix solutions were replaced in the populations of all treatments. This step was essential to ensure an oxygenated solution and to prevent infection from the solution mix that contained excess debris. In the HI treatment, resources were left for 2 h before removal to ensure full satiation.

The experiment was conducted over several blocks. Block 1 of the experiment was initiated in the climate rooms of the Department at the University where the experiments took place. However, due to the COVID-19 pandemic, experiments were forced to cease across the University. As such, block-1 treatments were transferred to a room on private premises 11 weeks after the initiation of the experiment and for the remainder of the experiment. Block 2 and block 3 were both initiated and completed within the same room at the same location. Temperature within the room was monitored three times a day (Fig. S1A) using a digital thermometer, which also recorded maximum and minimum daily temperature (Fig. S1B). This careful monitoring of working conditions during the pandemic disruption ensured that the temperatures were within the temperature range experienced within the climate room at the Departmental laboratory. Indeed, throughout the course of the experiment, temperature fluctuations never exceeded the maximum and minimum temperatures experienced within the climate room located at the Department (Fig. S1; grey horizontal lines indicate climate room temperature extremes).

**Image digitization**

To determine the impact of variation in resource availability on changes in body size of individuals within populations across treatments, we took photographs once a week of all populations. We placed a population on an A4 size light table and within an opened bottomed box (to allow light through). The top of the box had an 8cm diameter aperture to allow for a camera lens. A Canon EOS 700D digital SLR camera and a Canon 18-135mm EFS lens were attached to a tripod (Vanguard Alta Pro 263AP) so the lens was facing downwards and protruding into the top aperture of the box. The lens was set at a constant height of 29cm above the population. The camera settings were kept constant and set manually at F5.6, ISO 200 and a shutter speed of 1/100. All images were digitized and analysed using the CellProfiler software (Lamprecht *et al.* 2007). CellProfiler is a flexible open-source software allowing for user specific settings. Based on the user settings, the software identifies possible individuals within the image and calculates the relevant data (*e.g.*, area of an individual) (Fig. S2). Image settings were selected depending on the species in question, as there are significant size differences between *S. mediterranea* and *D. tahitiensis*. Specifically, the required minimum and maximum values of the parameters used to identify individuals in the image (the ‘object diameter identification’ parameter) differed between species. The specific user defined settings were then used for the relevant species. User defined settings were saved as a CellProfiler project file and is accessible to use via the online supplementary material files ‘Smed.cpproj’ and ‘Dtah.cpproj’.

In the case of *Dugesia tahitiensis*, the procedure differed slightly for week 19 for the high quality, seven day resource interval treatments for all resource intakes (high, standard and reduced intake). In these treatments high population numbers (>100 individuals per population) became problematic when trying to analyse size of individuals within some populations. In these populations the specific CellProfiler settings did not isolate all individuals within the image correctly; some individuals were either identified as two individuals or two individuals were identified as one if they were overlapping, resulting in the program not able to differentiate between them. The sensitivity of the settings within the program were adjusted to try and account for the higher number of individuals within a population. However, there were still large discrepancies between the number of individuals identified by the program and the count by hand. Given this issue, one quarter of the population/petri dish was sub-sampled to calculate the number of individuals and measure the body size of the individuals to use as a proxy measurement to extrapolate from. Here, we divided each population into quarters (*i.e.*, one vertical and one horizontal delineation of the population giving four equal sized parts). A quarter was then randomly chosen to analyse using the user defined settings within the CellProfiler project file (Dtah.cpproj). The results from the image analysis were then extrapolated for the given population by simply reproducing the data for the quadrants not measured. For both species, before an image was taken, if individuals were clumped together, we allowed individuals time to move around freely and become less aggregated. This ensured that when the photograph was taken individuals were more evenly distributed within the population. Finally, because the individual area data calculated by CellProfiler is provided in pixel values, we converted all values into mm^2^ by multiplying pixel values by a conversion factor. The scale for all images were identified by a marker that was present in all the petri dishes that housed the treatment populations. Given that camera settings, focal length, and height of the camera were the same for all images taken, we used the marker in the petri dishes as a scale. The length of the marker was measured as 4mm, this scale measurement was then used to calculate the equivalent length in pixels (101.47 px), from which we calculated the conversion factor. We divided the scale length by the pixel length to calculate the conversion factor (0.0394), the final step was to multiply all the area data in pixels by 0.0394^2^, thus providing area in mm^2^. Area data in mm^2^ was then used in all subsequent analysis.

**Statistical analysis**

*Size distribution*

To model the size spectra slope values (*b*) as a function of covariates, a Gaussian model was used (Eq. 2) with treatment (*T*, categorical with 12 levels), week (*W*, 19 weeks) and block(*B*, 3 blocks) as fixed variables, and population as a random effect. A Gaussian distribution was chosen over a Gamma distribution as the model with a Gaussian distribution was the best fit (Table S1).

${Slope value}_{i} \sim Normal(\mu_{i}, \sigma)$

$$\mu_{i}=\alpha_{T[i]}+\beta_{W*T\left[ i \right]} + \gamma_{B[i]}$$

$$\alpha_{T} \sim\mathrm{Normal}(x,z )$$

$\beta_{W} \sim\mathrm{Normal}(a,b )$

$$\gamma_{B} \sim\mathrm{Normal}\left( 0, 1 \right)$$

$\sigma\sim\mathrm{Exponential}\left( 1 \right)$	(2)

The model in Eq. (2) predicts the posterior mean size spectra slope value (*b*). We applied the model to the estimated mean *b* values and within the model we varied the mean (*x*) and standard deviation (*z*) in the α_T_ prior and the mean (*a*) and standard deviation (*b*) in the β_W_ prior as the expectation is shrinkage and regrowth of individuals across treatments. We then conducted a model comparison to ensure best model fit; models with the lowest WAIC values were taken as best fit (Table S2 & S3). In the case of *S. mediterranea* the model with the best fit had a mean value (*x*) of -0.75 and standard deviation value (*z*) of 1 for the *α_T_* prior, and mean (*a*) of 0 and standard deviation (*b*) of 0.05 for the β_W_ prior (Table S2). For *D. tahitiensis* model with the best fit had a mean value (*x*) of -0.61 and standard deviation value (*z*) of 0.5 for the *α_T_* prior, and mean (*a*) of 0 and standard deviation (*b*) of 1 for the β_W_ prior (Table S3).

*Population size*

*S. mediterranea*

Population counts were modelled using a Gaussian multilevel model and adaptive priors (Eq. 4), with treatment (*T*, categorical with 12 levels), week (*W*, 19 weeks) and block (*B*, 3 blocks) as fixed variables, and population (*P*, 60 populations) as a random effect. A Gaussian distribution was chosen over a Poisson distribution as the model with a Gaussian distribution was the best fit (Table S4).

${Counts}_{i} \sim Normal(\mu_{i}, \sigma)$

$$\mu_{i}=\alpha_{T[i]}+\beta_{W*T\left[ i \right]}+\beta_{{W\left[ i \right]}^{2}} + \gamma_{B[i]}$$

$$\alpha_{T} \sim\mathrm{Normal}(x, z)$$

$\beta_{W} \sim\mathrm{Normal}(a, b)$

$$\gamma_{B} \sim\mathrm{Normal}\left( 0, 1 \right)$$

$\sigma\sim\mathrm{Exponential}\left( 1 \right)$	(3)

The model in Eq. (3) predicts the posterior mean population. We applied the model to the population counts for all blocks and varied the mean (*x*) and standard deviation (*z*) in the α_T_ prior and the mean (*a*) and standard deviation (*b*) in the β_W_ prior. Following the model simulations based on the various mean and standard deviation values, we conducted a model comparison to ensure best model fit (models with the lowest WAIC values were taken as best fit, see Table S5). The model with the best fit had a mean value (*x*) of 23.43 and standard deviation value (*z*) of 1 for the *α_T_* prior, and mean (*a*) of 0 and standard deviation (*b*) of 1 for the β_W_ prior (Table S5).

*D. tahitiensis*

Population counts were modelled using a Poisson multilevel model with a log link function and adaptive priors (Eq. 4), with treatment (*T*, categorical with 12 levels), week (*W*, 19 weeks) and block (*B*, 3 blocks) as fixed variables, and population (*P*, 60 populations) as a random effect. A Poisson distribution was chosen over a Gaussian distribution as the model with a Poisson distribution was the best fit (Table S4).

${Counts}_{i} \sim Poisson(\mu_{i}, \sigma)$

$$\mu_{i}=\alpha_{T[i]}+\beta_{W*T\left[ i \right]}+\beta_{{W\left[ i \right]}^{2}} + \gamma_{B[i]}$$

$$\alpha_{T} \sim\mathrm{Normal}(x, z)$$

$\beta_{W} \sim\mathrm{Normal}(a, b)$

$$\gamma_{B} \sim\mathrm{Normal}\left( 0, 1 \right)$$

$\sigma\sim\mathrm{Exponential}\left( 1 \right)$	(4)

As with *S. mediterranea*, the model in Eq. (4) predicts the posterior mean population. We applied the model to the population counts for all blocks and varied the mean (*x*) and standard deviation (*z*) in the α_T_ prior and the mean (*a*) and standard deviation (*b*) in the β_W_ prior. Following the model simulations based on the various mean and standard deviation values, we conducted a model comparison to ensure best model fit (models with the lowest WAIC values were taken as best fit, see Table S6). The model with the best fit had a mean value (*x*) of 23.89 and standard deviation value (*z*) of 1 for the *α_T_* prior, and mean (*a*) of 0 and standard deviation (*b*) of 1 for the β_W_ prior (Table S6).

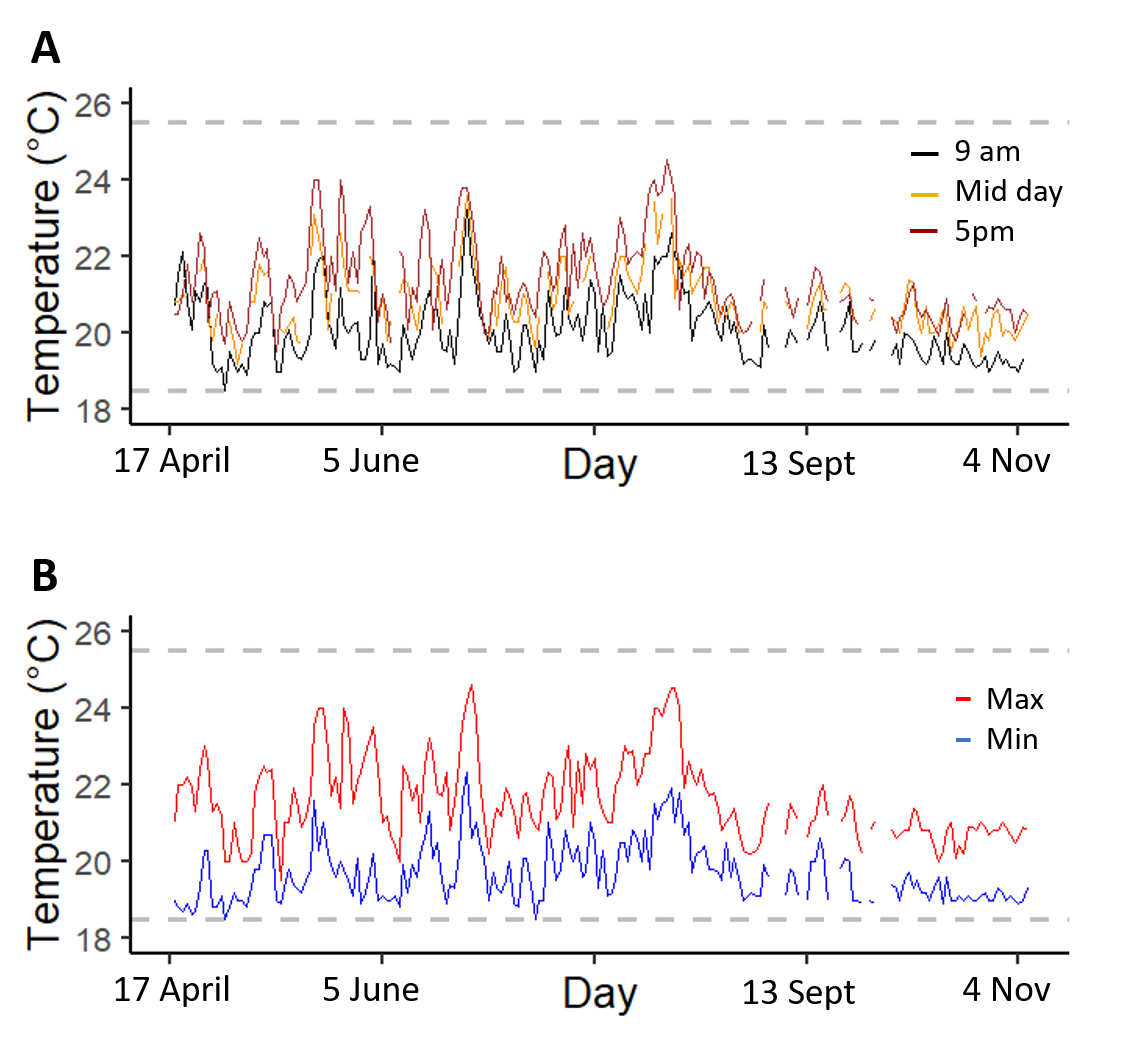

**Figure S1.** Temperature measurements within the private premises where the experiment was conducted from 17^th^ April 2020 until the end of the experiment on the 4^th^ of November 2020. A) Temperature readings at three time points throughout the day: 9am (black line), midday (orange line), 5pm (red line). B) Daily minimum (blue line) and maximum (red line) temperature over the time-period indicated. Grey dashed lines indicate minimum and maximum temperatures reached within the climate room in the laboratory based at the University where the experiments took place, where block 1 of our experiment was initiated, but from where we had to move the experiment elsewhere due to the pandemic lockdown.

**Table S1.** Model comparison output of a Gaussian vs. Gamma distribution for size slope spectra (*b*) analysis of both species. The distribution highlighted in bold indicates distribution of best fit used in the final analysis for both species (see Table S4 & S5). NA indicates model did not converge.

| **Model** | **Distribution** | **WAIC** | **SE** |
| --- | --- | --- | --- |
| ***S. mediterranea*** |  |  |  |
| *Slope value ~ Treatment * Week + Block + (1 \| population)* | Gamma | 2762.87 | 125.03 |
| *Slope value ~ Treatment + Week + Block + (1 \| population)* | Gamma | 2761.44 | 124.21 |
| *Slope value ~ Treatment + Week +* ${Week}^{2}$ *+ Block + (1 \| population)* | Gamma | 6632.55 | 30.44 |
| *Slope value ~ Treatment * Week + (1 \| population)* | Gamma | 2777.38 | 124.42 |
| *Slope value ~ Treatment + Week + (1 \| population)* | Gamma | 2773.01 | 124.01 |
| ***Slope value ~ Treatment * Week + Block + (1 \| population)*** | **Gaussian** | **2699.15** | **107.66** |
| ***Slope value ~ Treatment + Week + Block + (1 \| population)*** | **Gaussian** | **2711.41** | **107.96** |
| ***Slope value ~ Treatment + Week +*** $\boldsymbol{Week}^{\boldsymbol{2}}$ ***+ Block + (1 \| population)*** | **Gaussian** | **2711.09** | **107.50** |
| ***Slope value ~ Treatment * Week + (1 \| population)*** | **Gaussian** | **2697.92** | **106.83** |
| ***Slope value ~ Treatment + Week + (1 \| population)*** | **Gaussian** | **2708.67** | **107.65** |
| ***D. tahitiensis*** |  |  |  |
| *Slope value ~ Treatment * Week + Block + (1 \| population)* | Gamma | 2645.27 | 215.72 |
| *Slope value ~ Treatment + Week + Block + (1 \| population)* | Gamma | 2644.63 | 212.2 |
| *Slope value ~ Treatment + Week +* ${Week}^{2}$ *+ Block + (1 \| population)* | Gamma | NA | NA |
| *Slope value ~ Treatment * Week + (1 \| population)* | Gamma | 2656.65 | 208.46 |
| *Slope value ~ Treatment + Week + (1 \| population)* | Gamma | 2656.54 | 204.75 |
| ***Slope value ~ Treatment * Week + Block + (1 \| population)*** | **Gaussian** | **2417.63** | **153.08** |
| ***Slope value ~ Treatment + Week + Block + (1 \| population)*** | **Gaussian** | **2446.35** | **149.15** |
| ***Slope value ~ Treatment + Week +*** $\boldsymbol{Week}^{\boldsymbol{2}}$ ***+ Block + (1 \| population)*** | **Gaussian** | **2419.48** | **150.06** |
| ***Slope value ~ Treatment * Week + (1 \| population)*** | **Gaussian** | **2423.63** | **153.22** |
| ***Slope value ~ Treatment + Week + (1 \| population)*** | **Gaussian** | **2446.49** | **147.91** |

**Table S2.** Model comparison output for size slope spectra (*b*) analysis of *Schmidtea mediterranea*. The final model used in the analysis was the model with the lowest WAIC value (in bold).

| **Model** | **WAIC** | **SE** |
| --- | --- | --- |
| 1) ***S. mediterranea* priors for:** $\boldsymbol{\alpha\sim}\mathbf{Normal}\left( \boldsymbol{-0.75, 1} \right)$   $\boldsymbol{\beta}\boldsymbol{\sim}\mathbf{Normal}\left( \boldsymbol{0,0.5} \right)$ | **2696.56** | **107.46** |
| 2) *S. mediterranea* flat priors | 2697.12 | 106.85 |
| 3) *S. mediterranea* priors for: $\alpha\sim\mathrm{Normal}\left( -0.75, 0.5 \right)$  $\beta\sim\mathrm{Normal}\left( 0,0.5 \right)$ | 2698.59 | 107.50 |
| 4) *S. mediterranea* priors for: $\alpha\sim\mathrm{Normal}\left( -0.75, 1 \right)$  $\beta\sim\mathrm{Normal}\left( 0,1 \right)$ | 2698.78 | 107.26 |
| 5) *S. mediterranea* priors for: $\alpha\sim\mathrm{Normal}\left( -0.75, 0.5 \right)$  $\beta\sim\mathrm{Normal}\left( 0,1 \right)$ | 2700.70 | 107.33 |

**Table S3.** Model comparison output for size slope spectra (*b*) analysis of *Dugesia tahitiensis*. The final model used in the analysis was the model with the lowest WAIC value (in bold).

| **Model** | **WAIC** | **SE** |
| --- | --- | --- |
| 1) ***D. tahitiensis* priors for:** $\boldsymbol{\alpha\sim}\mathbf{Normal}\left( \boldsymbol{-0.61, 0.5} \right)$   $\boldsymbol{\beta}\boldsymbol{\sim}\mathbf{Normal}\left( \boldsymbol{0,1} \right)$ | **2419.05** | **150.08** |
| 2) *D. tahitiensis* priors for: $\alpha\sim\mathrm{Normal}\left( -0.61, 1 \right)$  $\beta\sim\mathrm{Normal}\left( 0, 1 \right)$ | 2419.76 | 150.05 |
| 3) *D. tahitiensis* priors for: $\alpha\sim\mathrm{Normal}\left( -0.61, 0.5 \right)$  $\beta\sim\mathrm{Normal}\left( 0, 0.5 \right)$ | 2419.77 | 150.13 |
| 4) *D. tahitiensis* flat priors: | 2419.80 | 149.97 |
| 5) *D. tahitiensis* priors for: $\alpha\sim\mathrm{Normal}\left( -0.61, 1 \right)$  $\beta\sim\mathrm{Normal}\left( 0, 0.5 \right)$ | 2424.71 | 151.57 |

**Table S4.** Model comparison output of a Gaussian vs. Poisson distribution for the population count analysis of both species. The distribution highlighted in bold indicates distribution of best fit used in the final analysis for both species. NA indicates model did not converge.

| **Model** | **Distribution** | **WAIC** | **SE** |
| --- | --- | --- | --- |
| ***S. mediterranea*** |  |  |  |
| *Counts ~ Treatment * Week + Block + (1 \| population)* | Poisson | 6183.39 | 54.32 |
| *Counts ~ Treatment + Week + Block + (1 \| population)* | Poisson | 6184.11 | 54.94 |
| *Counts ~ Treatment + Week +* ${Week}^{2}$ *+ Block + (1 \| population)* | Poisson | NA | NA |
| *Counts ~ Treatment * Week + (1 \| population)* | Poisson | 6674.96 | 76.67 |
| *Counts ~ Treatment + Week + (1 \| population)* | Poisson | 6677.61 | 77.51 |
| ***Counts ~ Treatment * Week + Block + (1 \| population)*** | **Gaussian** | **6161.11** | **65.76** |
| ***Counts ~ Treatment + Week + Block + (1 \| population)*** | **Gaussian** | **6160.14** | **66.06** |
| *Counts ~ Treatment + Week +* ${Week}^{2}$ *+ Block + (1 \| population)* | Gaussian | 6328.16 | 75.18 |
| *Counts ~ Treatment * Week + (1 \| population)* | Gaussian | 6618.71 | 69.82 |
| *Counts ~ Treatment + Week + (1 \| population)* | Gaussian | 6621.91 | 70.06 |
| ***D. tahitiensis*** |  |  |  |
| ***Counts ~ Treatment * Week + Block + (1 \| population)*** | **Poisson** | **6424.94** | **38.49** |
| ***Counts ~ Treatment + Week + Block + (1 \| population)*** | **Poisson** | **6425.26** | **38.45** |
| *Counts ~ Treatment + Week +* ${Week}^{2}$ *+ Block + (1 \| population)* | Poisson | 7700.00 | 81.39 |
| *Counts ~ Treatment * Week + (1 \| population)* | Poisson | 7700.15 | 81.24 |
| ***Counts ~ Treatment + Week + (1 \| population)*** | **Poisson** | **6293.64** | **29.91** |
| *Counts ~ Treatment * Week + Block + (1 \| population)* | Gaussian | 7221.58 | 93.01 |
| *Counts ~ Treatment + Week + Block + (1 \| population)* | Gaussian | 7222.3 | 92.44 |
| *Counts ~ Treatment + Week +* ${Week}^{2}$ *+ Block + (1 \| population)* | Gaussian | 8977.67 | 72.56 |
| *Counts ~ Treatment * Week + (1 \| population)* | Gaussian | 8983.11 | 72.21 |
| *Counts ~ Treatment + Week + (1 \| population)* | Gaussian | 7225.23 | 91.14 |

**Table S5.** Model comparison output for the population count analysis of *Schmidtea mediterranea*. The final model used in the analysis was the model with the lowest WAIC value (in bold).

| **Model** | **WAIC** | **SE** |
| --- | --- | --- |
| 2) ***S. mediterranea* priors for:** $\boldsymbol{\alpha\sim}\mathbf{Normal}\left( \boldsymbol{23.43, 1} \right)$   $\boldsymbol{\beta}\boldsymbol{\sim}\mathbf{Normal}\left( \boldsymbol{0,1} \right)$ | **5667.20** | **68.11** |
| 2) *S. mediterranea* flat priors for the average intercept | 5668.45 | 67.74 |
| 3) *S. mediterranea* priors for: $\alpha\sim\mathrm{Normal}\left( 23.43, 1 \right)$  $\beta\sim\mathrm{Normal}\left( 0,0.5 \right)$ | 5669.90 | 67.41 |
| 4) *S. mediterranea* priors for: $\alpha\sim\mathrm{Normal}\left( 23.43, 0.5 \right)$  $\beta\sim\mathrm{Normal}\left( 0,1 \right)$ | 5670.26 | 67.95 |
| 5) *S. mediterranea* priors for: $\alpha\sim\mathrm{Normal}\left( 23.43, 0.5 \right)$  $\beta\sim\mathrm{Normal}\left( 0,1 \right)$ | 5672.36 | 68.26 |

**Table S6.** Model comparison output for the population count analysis of *Dugesia tahitiensis*. The final model used in the analysis was the model with the lowest WAIC value (in bold).

| **Model** | **WAIC** | **SE** |
| --- | --- | --- |
| 1) ***D. tahitiensis* priors for:** $\boldsymbol{\alpha\sim}\mathbf{Normal}\left( \boldsymbol{23.89, 1} \right)$   $\boldsymbol{\beta}\boldsymbol{\sim}\mathbf{Normal}\left( \boldsymbol{0,1} \right)$ | **7222.06** | **91.35** |
| 2) *D. tahitiensis* flat priors for the average intercept | 7224.22 | 92.51 |
| 3) *D. tahitiensis* priors for: $\alpha\sim\mathrm{Normal}\left( 23.89, 0.5 \right)$  $\beta\sim\mathrm{Normal}\left( 0,1 \right)$ | 7227.17 | 92.80 |
| 4) *D. tahitiensis* priors for: $\alpha\sim\mathrm{Normal}\left( 23.43, 1 \right)$  $\beta\sim\mathrm{Normal}\left( 0,0.5 \right)$ | 7229.32 | 92.99 |
| 5) *D. tahitiensis* priors for: $\alpha\sim\mathrm{Normal}\left( 23.43, 0.5 \right)$  $\beta\sim\mathrm{Normal}\left( 0,0.5 \right)$ | 7232.20 | 93.85 |

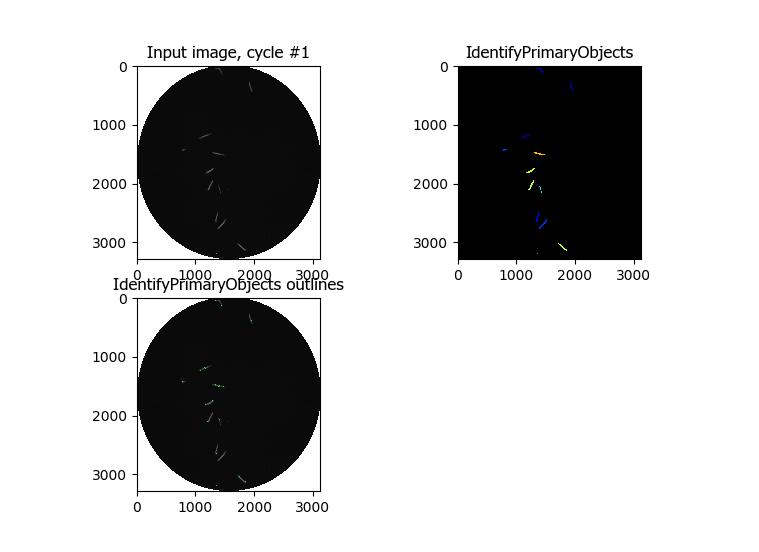

**Figure S2.** Example of how the CellProfiler program identifies individuals within each population of the experiment. The program converts an image to black and white (top left, ‘Input image’), identifies the outlines of objects in the image and sets the outlines that conform to the user defined settings (bottom left, ‘IdentifyPrimaryObjects outlines’). The relevant outlines are then set as primary objects for subsequent analysis (top right, ‘IdentifyPrimaryObjects’). The identified areas are then measured.

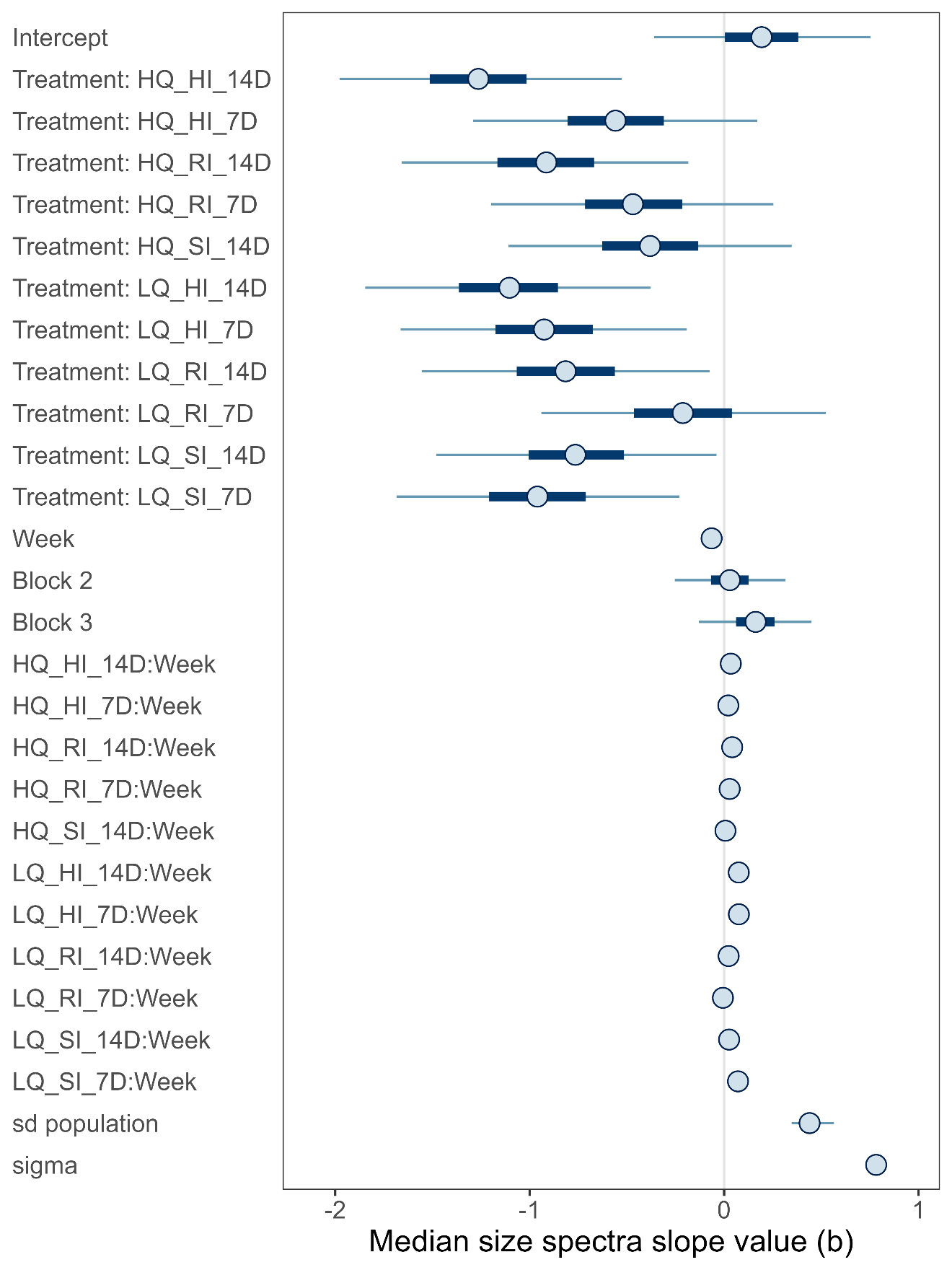

**Figure S3.** Median estimated size spectra slope values from the Bayesian model for *Schmidtea mediterranea*. Shown are posterior median values with 95% (thin lines) and 50% (thick lines) compatibility intervals. Also shown is the standard deviation for the distribution of intercepts across the different populations (random effect) (sd_pop_id_Intercept) and the global variation (sigma). HQ – high quality, LQ – low quality; HI – high intake, SI – standard intake, RI – reduced intake; 7D – seven day feeding interval, 14D – 14 day feeding interval.

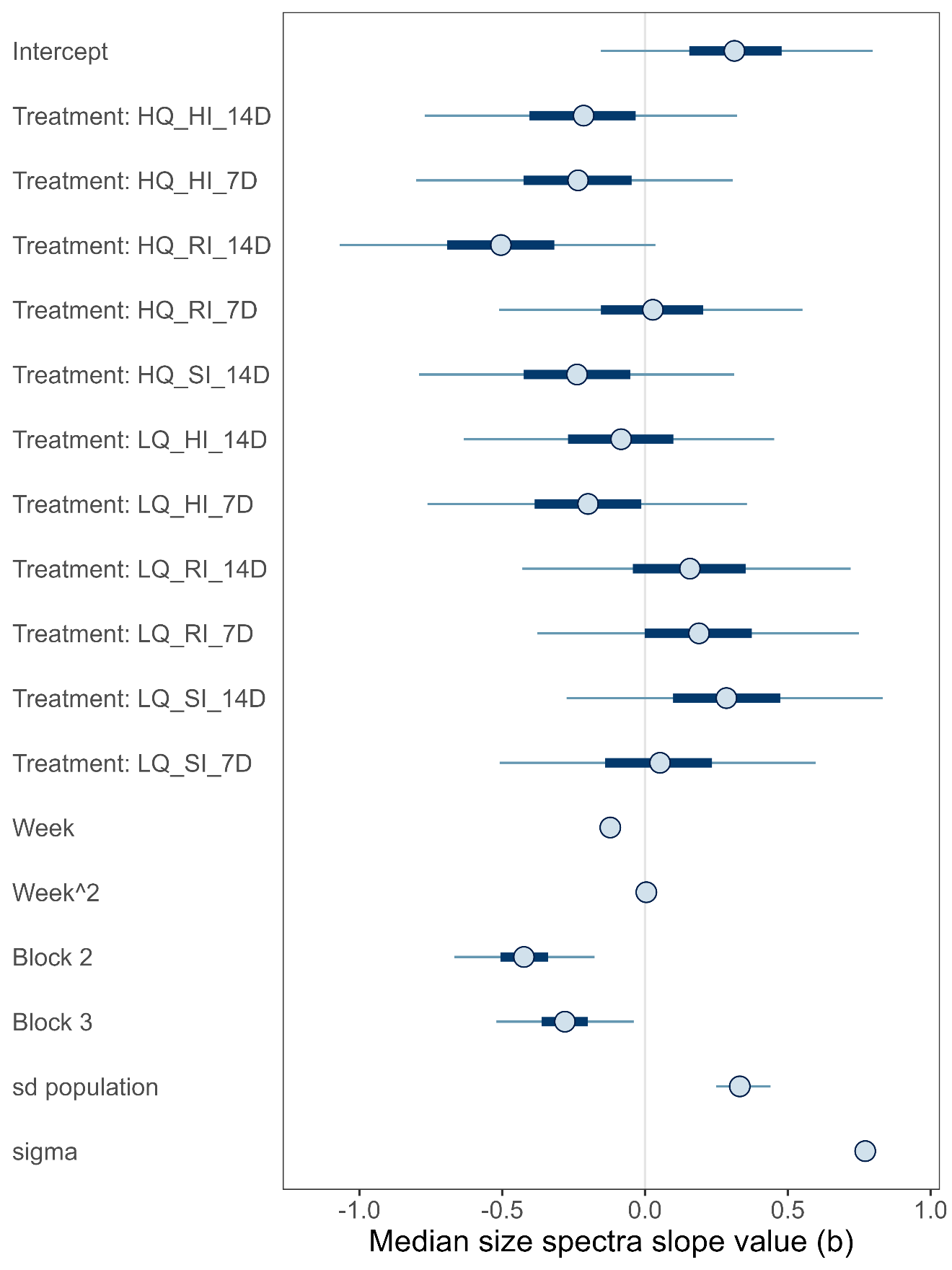

**Figure S4.** Mean estimated size spectra slope values from the Bayesian model for *Dugesia tahitiensis*. Shown are posterior median values with 95% (thin lines) and 50% (thick lines) compatibility intervals. Also shown is the standard deviation for the distribution of intercepts across the different populations (random effect) (sd_pop_id_Intercept) and the global variation (sigma). HQ – high quality, LQ – low quality; HI – high intake, SI – standard intake, RI – reduced intake; 7D – seven day feeding interval, 14D – 14 day feeding interval.

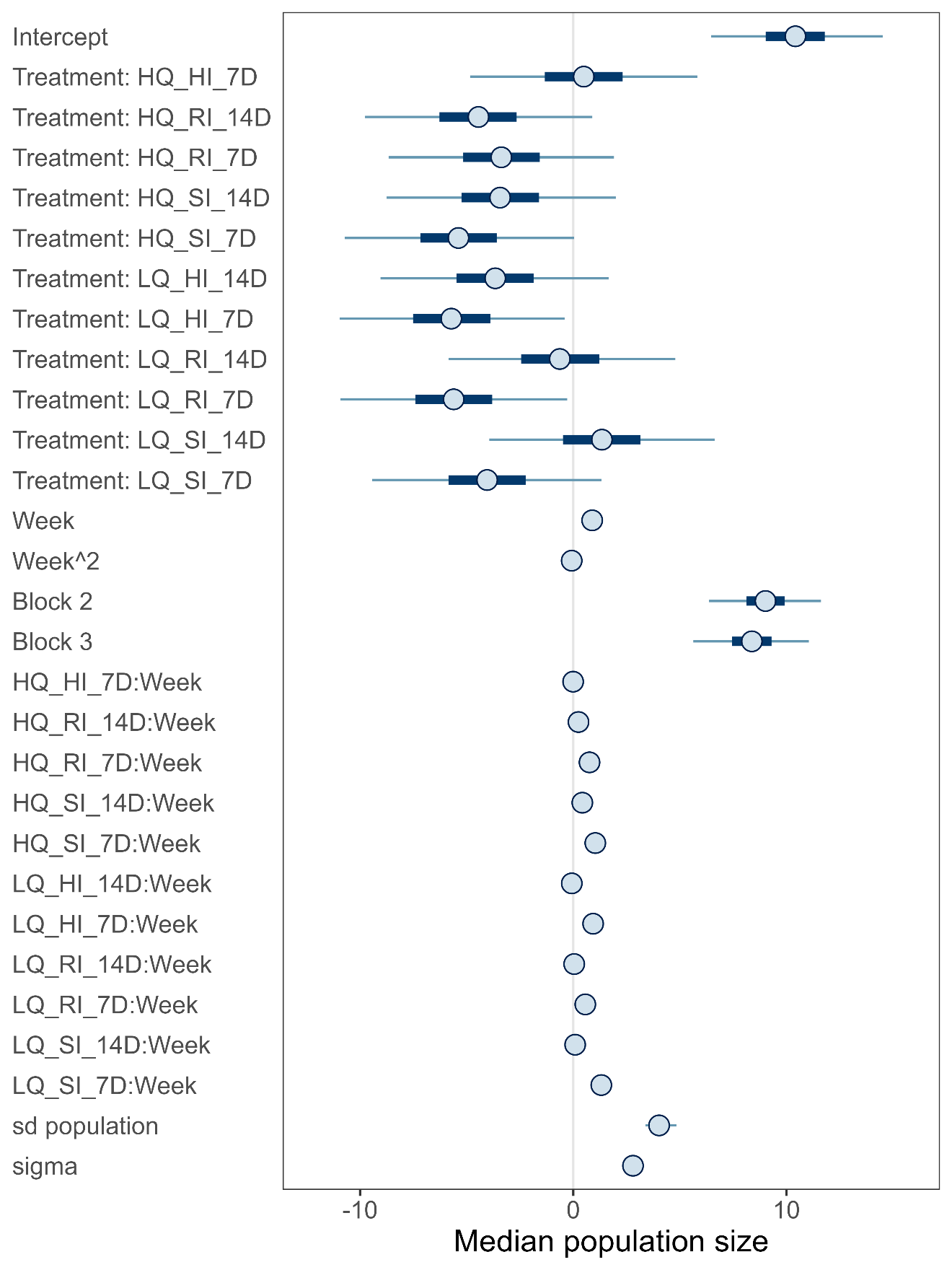

**Figure S5.** Median estimated count values from the Bayesian model for *Schmidtea mediterranea*. Shown are posterior median values with 95% (thin lines) and 50% (thick lines) compatibility intervals. Also shown is the standard deviation for the distribution of intercepts across the different populations (random effect) (sd_pop_id_Intercept) and the global variation (sigma). HQ – high quality, LQ – low quality; HI – high intake, SI – standard intake, RI – reduced intake; 7D – seven day feeding interval, 14D – 14 day feeding interval.

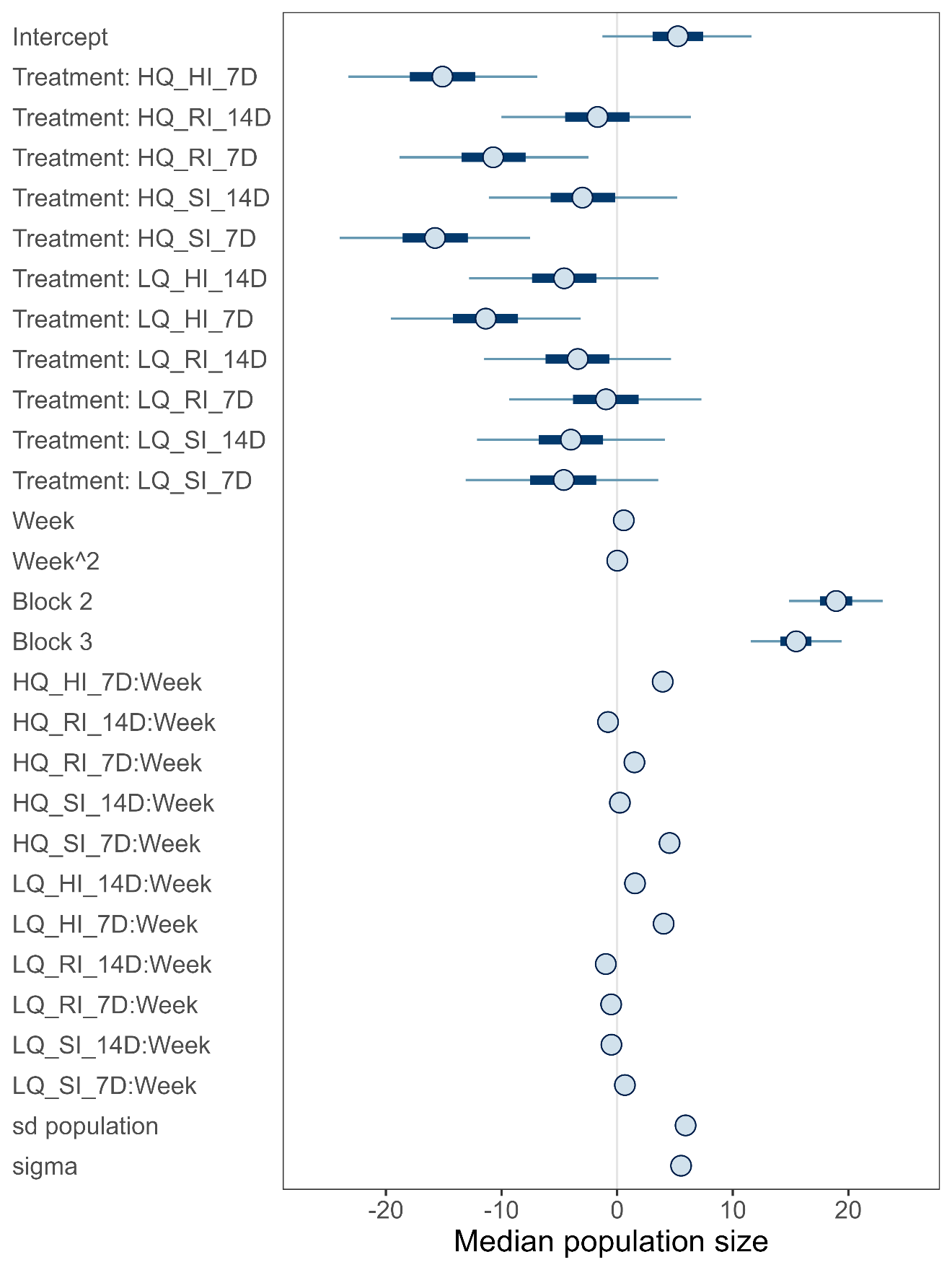

**Figure S6.** Mean estimated count values from the Bayesian model for *Dugesia tahitiensis*. Shown are posterior median values with 95% (thin lines) and 50% (thick lines) compatibility intervals. Also shown is the standard deviation for the distribution of intercepts across the different populations (random effect) (sd_pop_id_Intercept) and the global variation (sigma). HQ – high quality, LQ – low quality; HI – high intake, SI – standard intake, RI – reduced intake; 7D – seven day feeding interval, 14D – 14 day feeding interval.

*References*

Elliott, S.A. & Sánchez Alvarado, A. (2013). The history and enduring contributions of planarians to the study of animal regeneration. *Wiley Interdiscip. Rev. Dev. Biol.*, 2, 301–326.

Lamprecht, M.R., Sabatini, D.M. & Carpenter, A.E. (2007). CellProfiler^TM^: free, versatile software for automated biological image analysis. *BioTechniques*, 42, 71–75.

Sousa, N. de & Adell, T. (2018). Maintenance of Schmidtea mediterranea in the Laboratory. *Bio-Protoc.*, 8, e3040.
